# The fate of secretory cells during intestinal homeostasis, regeneration, and tumor formation is regulated by Tcf4

**DOI:** 10.1101/2024.07.11.603019

**Authors:** Lucie Janeckova, Monika Stastna, Dusan Hrckulak, Linda Berkova, Jan Kubovciak, Jakub Onhajzer, Vitezslav Kriz, Stela Dostalikova, Tereza Mullerova, Katerina Vecerkova, Marketa Tenglerova, Stepan Coufal, Klara Kostovcikova, Richard S Blumberg, Dominik Filipp, Konrad Basler, Tomas Valenta, Michal Kolar, Vladimir Korinek

**Author notes:** Corresponding Authors: *Vladimir Korinek and Lucie Janeckova Laboratory of Cell and Developmental Biology Institute of Molecular Genetics, Czech Academy of Sciences Videnska 1083 Prague 4, 142 20, Czech Republic Email:**, Email:.

## Abstract

The single-layer epithelium of the gastrointestinal tract is a dynamically renewing tissue that ensures nutrient absorption, secretory and barrier functions and is involved in immune responses. The basis for this homeostatic renewal is the Wnt signaling pathway. Blocking this pathway can lead to epithelial damage, while its abnormal activation can result in the development of intestinal tumors.

In this study, we investigated the dynamics of intestinal epithelial cells and tumorigenesis using a conditional mouse model. Using single-cell and bulk RNA sequencing and histological analysis, we elucidated the cellular responses following the loss of specific cell types. We focused on the fate of cells in the lower parts of the intestinal crypts and the development of colon adenomas. By partially inactivating the transcription factor Tcf4, a key effector of the Wnt signaling pathway, we analyzed the regeneration of isolated hyperproliferative foci (crypts). Our results suggest that the damaged epithelium is not restored by a specific regeneration program associated with oncofetal gene production, but rather by a standard homeostatic renewal pathway. Moreover, disruption of Tcf4 in secretory progenitors resulted in a significant shift in the cell lineage from Paneth cells to goblet cells, characterized by morphological changes and loss of Paneth cell-specific genes. We also found that hyperactivation of the Wnt signaling pathway in colonic adenomas correlated with the upregulation of genes typical of Paneth cells in the intestine, followed by the emergence of secretory tumor cells producing the Wnt3 ligand. The absence of Tcf4 led to a phenotypic shift of the tumor cells towards goblet cells.

Our study presents a new model of epithelial regeneration based on the genetically driven partial elimination of intestinal crypts. We highlight the critical role of Tcf4 in the control of cell lineage decisions in the intestinal epithelium and colon tumors.

**GRAPHICAL ABSTRACT:** 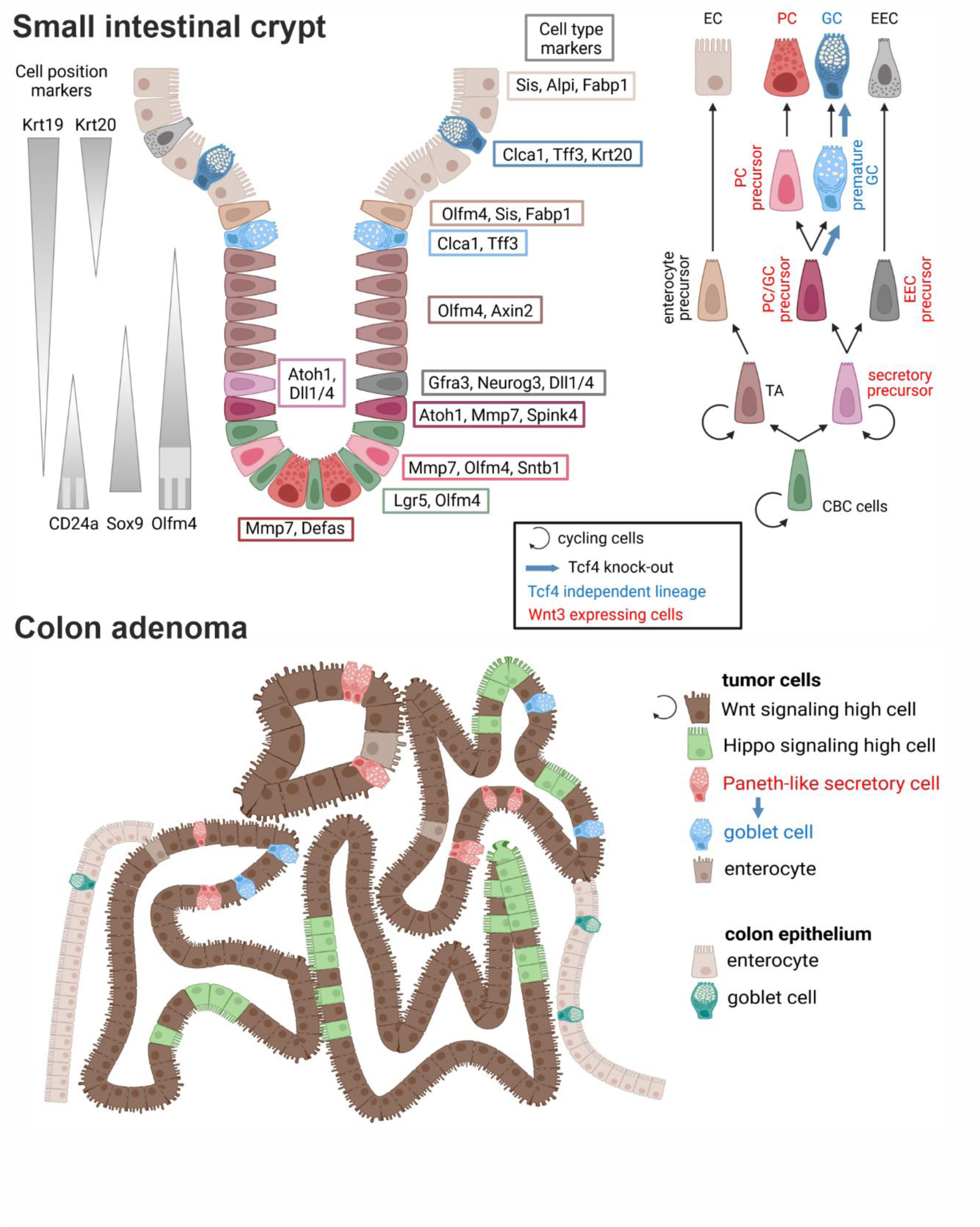

## INTRODUCTION

The small intestinal epithelium is a highly organized and dynamic structure that plays a crucial role in nutrient absorption and barrier function. Morphologically, it is characterized by its villi and crypts. The villi, finger-like protrusions that increase the surface area for absorption, are lined with a single layer of columnar epithelial cells. Beneath the villi lie the crypts of Lieberkühn, which harbor intestinal stem cells (ISCs) at the base of the crypts. These ISCs play a central role in the continuous renewal of the epithelium by giving rise to rapidly dividing progenitor cells that differentiate as they migrate upwards to the tips of the villi. The cell lineage decision takes place in the so-called differentiation zone, which is located at the +4/+5 position of the crypt base (1).

Goblet cells, the most common secretory cells in the small and large intestine, originate at the crypt base and migrate to the villi in the small intestine and to the luminal surface in the large intestine. There, they produce mucin 2 (Muc2), an important component of the protective mucus layer (2). The second most abundant secretory cells are Paneth cells, which are located at the bottom of the crypt. Paneth cells differ from goblet cells in their active Wnt signaling and create a niche for ISCs through antimicrobial protection and secretion of Wnt and Notch ligands (3). In colonic crypts, deep crypt secretory (DCS) cells, characterized by their expression of regenerating islet family member 4 (Reg4), are involved in the maintenance of the ISC niche (4). Although DCS cells do not produce Wnt ligands or lysozyme-containing granules like their counterparts in the small intestine, they share similarities with Paneth cells, and are therefore referred to as Paneth-like cells. Similar to Paneth cells in the small intestine, DCS cells are inserted between colonic stem cells and produce peptides and mucins for the host defense (5).

Both Paneth cells and goblet cells play an essential role in establishing a barrier between the host tissue and the intestinal microbiota residing in the intestinal lumen. Dysregulation of their functions is associated with the development of inflammatory bowel diseases such as Crohn’s disease and ulcerative colitis (6–8). The functionality of Paneth cells is closely linked to the activity of T cell factor 4 (Tcf4), which is encoded by the Tcf7l2 gene (9, 10). Mutations in TCF7L2 are associated with Crohn’s disease in the ileum in humans, as the depletion of Paneth cells leads to a reduced concentration of antimicrobial peptides alpha-defensins 5 and 6 (DEFA5/6), which results in chronic inflammation of the intestinal mucosa (9). Tcf4 acts as a downstream mediator of the Wnt signaling pathway and regulates gene expression in a Wnt-dependent manner. Binding of Wnt ligands to their receptors frizzled (FZD) on the surface of signal-receiving cells initiates a series of events that ultimately lead to translocation of transcriptional activator β-catenin into the nucleus. In the nucleus, β-catenin forms a complex with TCF/lymphoid enhancer factor (LEF) transcription factors and activates Wnt target genes (11, 12). Wnt signaling is particularly active in the lower regions of the intestinal crypts and promotes the proliferation and pluripotency of ISCs (13) as well as the maturation of Paneth cells (14).

While physiological levels of Wnt signaling are essential for cell proliferation and epithelial renewal, its aberrant activation can lead to uncontrolled stem cell proliferation, which is closely associated with tumorigenesis in the gastrointestinal tract (15, 16). Colorectal cancer (CRC) is characterized by progressive accumulation of genetic mutations that transform normal colonic epithelial cells into malignant tumors. One of the earliest and best documented events in colorectal carcinogenesis is inactivation of the adenomatous polyposis coli (*APC*) gene. This gene encodes a scaffold protein that is critical for the formation of the cytoplasmic β-catenin degradation complex. Consequently, mutations that inactivate APC lead to the accumulation of β-catenin, independent of external Wnt signaling, resulting in abnormal activation of the Wnt pathway (17).

The crucial role of Wnt/Tcf4 signaling for the function of the intestinal epithelium was underlined by studies with knockout mice. The complete knockout of Tcf7l2 in mice leads to perinatal lethality, which is mainly due to the lack of proliferative compartments in the small intestine (13). In addition, tissue-specific deletion of Tcf4 from the intestinal epithelium of adult mice leads to the absence of proliferating cells, and thus blocks the self-renewal capacity of the small intestinal epithelium (10, 18).

In this study, we performed a detailed analysis using single-cell and bulk RNA sequencing to investigate the effects of loss of factor Tcf4 on the fate of cells in the lower part of the crypt epithelium of the mouse small intestine. In addition to the conditional *Tcf7l2* gene alleles and the Cre drivers *Villin-CreERT2* and *Defa6-iCre,* we also used reporter mice *Rosa26-tdTomato* and *Mki67^RFP^*. The latter allowed us to follow the fate of Tcf4-deficient and dividing cells, respectively. The resulting phenotypes of Tcf4-deficient cells were compared to cells with intact Tcf4. Partial loss of Tcf4 in the stem cell compartment resulted in epithelial damage and formation of rapidly dividing regenerative crypts. Our results suggest that epithelial regeneration after the loss of Tcf4-deficient cells does not occur via a specific regenerative pathway, as might be expected, but rather as part of a "normal" homeostatic renewal. The loss of Tcf4 in the secretory cell compartment directs the differentiation of progenitor cells towards goblet cells. In our previous study, we observed significant upregulation of Paneth cell-specific gene expression in the mouse colonic epithelium within a few days after conditional *Apc* knockout (19). This finding allowed us to use the *Defa6-iCre* driver, which is active in the lineage of Paneth cells and their progenitors, to label tumor cells. In combination with the reporter allele, we were thus able to follow the developmental relationships (trajectories) within the tumor cells. We also observed that inactivation of the *Tcf7l2* gene redirected the tumor cells into the goblet cell lines.

## RESULTS

### Intestinal epithelial cell dynamics and microbiota composition are altered by inactivation of Tcf4 in a conditional *Tcf7l2^flox/flox^/Villin-CreER^T2^* mouse model

In our previous work, we have shown that inactivation of Tcf4 in all epithelial cells of the small intestine leads to the loss of proliferating cells and subsequent death of experimental animals between 8-16 days after tamoxifen administration in *Tcf7l2^flox/flox^/Villin-CreER^T2^*animals (18). However, since the loss of the *Tcf7l2* gene is not complete (100%), proliferating hyperplastic lesions are observed in the small intestine (Fig. 1A and Supplementary Fig. S1A). The absence of Tcf4 in the epithelium is consistent with the phenotype of reduced and misplaced lysozyme-positive, presumably Paneth cells, as reported by van Es and colleagues (10). We also observed an increased presence of regenerating islet-derived 3 beta (Reg3b), which in a healthy gut is primarily present at the interface between the crypts and the gut surface and was almost ubiquitously expressed in the Tcf4-deficient gut, including the remaining crypts that regenerate the epithelium (Fig. 1B). The expression of Reg3b is regulated by the intestinal microbiota and serves to protect the intestinal epithelium from inflammation (20, 21). Interestingly, when the mice were treated with antibiotic vancomycin in the drinking water one week before tamoxifen administration, i.e., Cre-mediated recombination of the floxed alleles, the epithelium, which previously had only a few foci of proliferating cells, was completely restored and the mortality of the mice was significantly reduced (Fig. 1C). We therefore investigated the composition of the gut microbiome in the small and large intestine of these mice. The gut of the wild-type (wt) animals showed a greater diversity of bacterial species and a higher abundance of probiotic strains such as *Lactobacillus*, *Bifidobacterium* (22) and *Dubosiella* (23). In contrast, the *Tcf7l2* conditional knockout (cKO) mice harbored a substantial concentration of bacterial strains, including *Enterococcus* and *Escherichia-Shigella*, which are commonly found in patients with inflammatory bowel disease (IBD) (24) (Fig. 1D).

**Figure 1.**
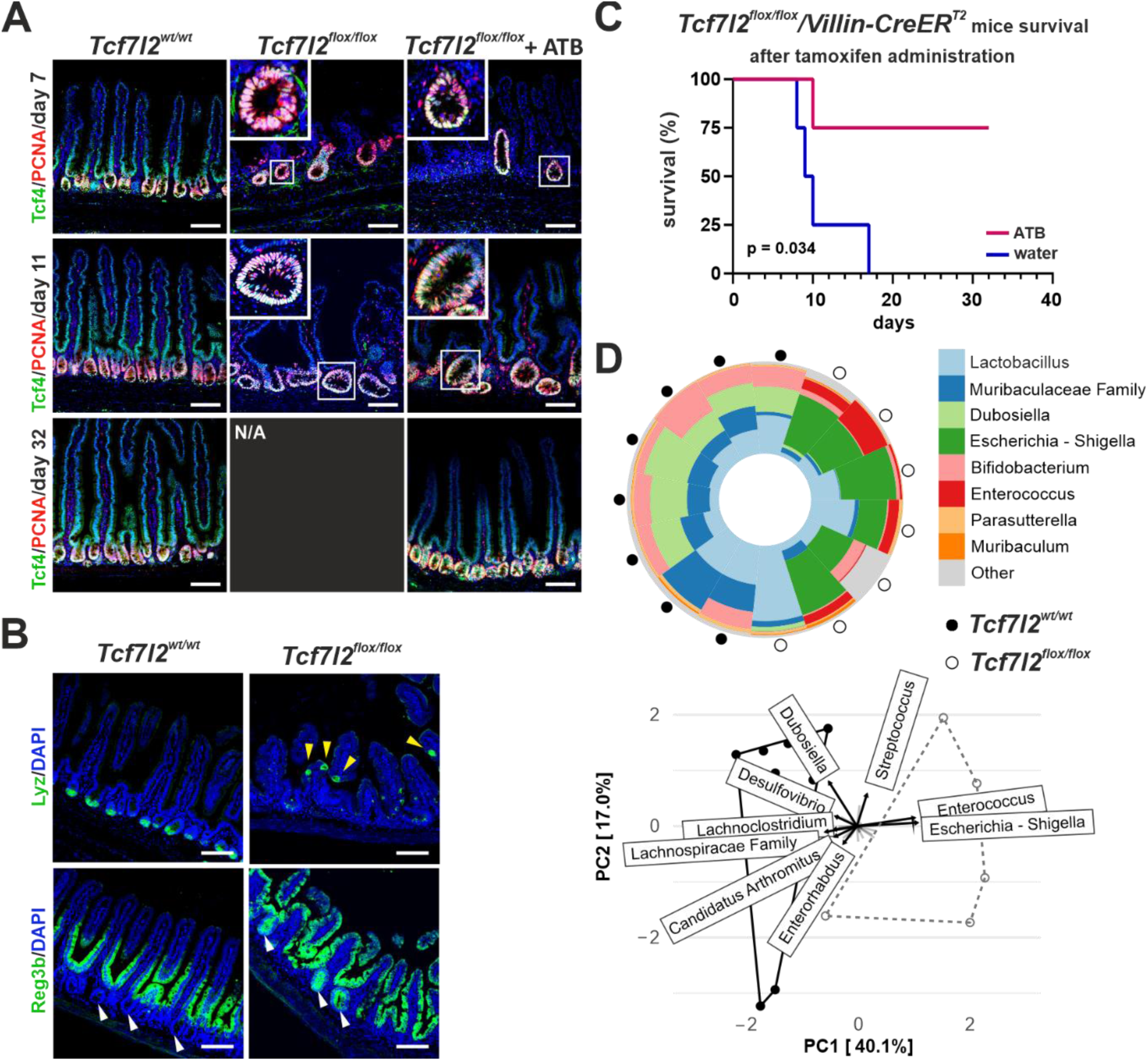
*Tcf7l2* knockout effect on intestinal epithelial regeneration and microbiota dynamics. A) Fluorescence micrographs of T-cell factor 4 (Tcf4, green) and proliferating cell nuclear antigen (PCNA, red) protein localization in the small intestine of wild-type *Tcf7l2^wt/wt^/VillinCreER^T2^*(*Tcf7l2^wt/wt^*) and knockout *Tcf7l2^flox/flox^ /VillinCreER^T2^* (*Tcf7l2^flox/flox^*) mice, with antibiotics (ATB) and without antibiotic treatment, over time after tamoxifen administration. The images show disruptions in the epithelial architecture of Tcf4-deficient mice that are partially restored by antibiotic treatment. The insets show increased magnification of the framed areas. Scale bar: 50 μm. B) Histological sections show the mislocalization and expression changes of the Paneth cell markers lysozyme (Lyz) and regenerating islet-derived 3 beta (Reg3b) seven days after tamoxifen administration. Yellow arrowheads indicate Lyz-positive cells on the villi; the white arrowheads in the lower left image show that the crypt bases in Tcf4^+^ epithelium are not positive for Reg3b, whereas they produce Reg3b in Tcf4-deficient epithelium (white arrowheads in the lower right figure). Scale bar: 50 μm. C) Kaplan-Meier survival curves for *Tcf7l2* knockout mice treated with either antibiotics or water after tamoxifen induction. Statistical significance (p = 0.034) shows that antibiotic treatment increases survival. D) Analysis of the composition of the microbiome in the distal small intestine of *Tcf7l2^wt/wt^* and *Tcf7l2^flox/flox^* mice. The pie chart shows the bacterial distribution in Tcf4-proficient mice (a solid black point on the outer perimeter of the chart) and Tcf4-deficient mice (an empty black ring). The principal component analysis (PCA) below the diagram illustrates the variance between bacterial populations, indicating shifts in the microbiota associated with the *Tcf7l2* status. Tcf7l2, transcription factor 7 like 2.

We then analyzed the phenotypes generated after crossing the cKO *Tcf7l2* allele with the *Villin-CreER^T2^* driver by transcriptional profiling. We investigated the cellular expression program of the regenerating parts of the intestine, i.e., hyperplastic crypts, compared to the proliferating cells of crypts with intact *Tcf7l2* gene. Therefore, we crossed *Tcf7l2^flox/flox^*/*Villin-CreER^T2^*mice in the homozygous state with the *Mki67^RFP^* strain, which produces the TagRFP red fluorescent protein in frame at the C-terminus of Ki67 (25). We then performed bulk RNA sequencing (RNA-seq) and single-cell (sc) RNA-seq analyses of dividing, i.e., Mki67-RFP-positive, cells sorted from hyperproliferative epithelial cells in which the *Tcf7l2* gene remained intact. The RNA expression profile of cells isolated from mice without *CreER^T2^* cassette served as a control. The resulting bulk RNA-seq analysis showed that the dividing cells derived from the hyperproliferative parts of the epithelium had a similar expression profile to the dividing cells localized in normal crypts. This comparison revealed (only) 18 significantly increased genes encoding proteins included in Gene Set Enrichment Analysis (GSEA) sets associated with cell growth and division (or metabolic activities). Among the 12 down-regulated genes, there were in total five genes encoding different α-defensins (Supplementary Fig. S1BC and Supplementary Table S2).

Interestingly, the scRNA-seq analysis showed a similar distribution of dividing cells in the regenerating epithelium of the small intestine and in the crypts of the control tissue. The results were the same regardless of whether antibiotics were administered or not (analysis without antibiotic treatment is shown only). Since we intentionally did not correct for the cell cycle, some cell populations "secondarily" split into several clusters depending on the stage of the cell cycle (Fig. 2A; heatmap visualization is shown in Supplementary Fig. S2). The analysis also had to take into account that the TagRFP protein is relatively stable and some TagRFP-positive cells may no longer be active in the cell cycle. In any case, we observed three clusters of cells that are positive for the stem cell marker leucine-rich repeat-containing G protein-coupled receptor 5 (*Lgr5*) (26) and olfactomedin 4 (*Olfm4*) (27). The first two (1 and 7) represent "classical" crypt base columnar (CBC) stem cells in phases G1/S and G2/M, respectively. The third cluster, number 14 (Lyz1^+^/Olfm4^+^ cells in Fig. 2B) is reminiscent of label-retaining cells (LRCs) that co-express stem cells [*Olfm4*, sex-determining region Y (SRY)-box 9 (*Sox9*)] and Paneth cell markers [defensins, *Reg4*, lysozyme 1 (*Lyz1*), matrix metallopeptidase 7 (*Mmp7*)] (28). Four clusters (2, 4, 5 and 6) represented transit amplifying (TA) cells; the cells in these and stem cell clusters had the highest levels of cell proliferation nuclear antigen (*PCNA*) and/or *Mki67*. Three clusters (3, 10, 11) expressed the regulator of secretory cell fate, atonal bHLH transcription factor 1 (*Atoh1*) (29), serine peptidase inhibitor, Kazal type 4 (*Spink4*) and the Atoh1 target gene, SAM pointed domain containing ETS transcription factor (*Spdef*) (30). Prediction of future gene expression of cells in clusters based on the ratio of spliced and unspliced mRNA (scRNA velocity) (31) showed a transition from cells in cluster 3, which may represent the earliest secretory progenitor cells, to clusters 10 and 11, which represent slightly more developed cells of the enteroendocrine [positive for neurogenin 3 (*Neurog3*) (32) and glial cell line derived neurotrophic factor family receptor alpha 3 (*Gfra3*)] (33) and goblet/Paneth cells [markers anterior gradient 2 (*Agr2*), trefoil factor 3 (*Tff3*), *Muc2* for goblet and *Lyz1*, *Defa17/24*, *Mmp7* for Paneth cells] (34, 35), respectively. Interestingly, secretory progenitor cells in particular were less abundant in the *Tcf7l2^flox/flox^* sample (Fig. 2A). Three clusters, 12, 8 and 13, represented the pathway to absorptive enterocytes [intestinal alkaline phosphatase (*Alpi*)-, sucrase-isomaltase (*Sis*)- and fatty acid binding protein 1 (*Fabp1*)-positive] (35, 36). The cells in cluster 9 were *Olfm4*- and *Lgr4*-positive and produced genes related to Paneth and goblet cells. However, the expression profile of these apparently secretory progenitor cells was very heterogeneous and some of these cells expressed enterocytic markers (Supplementary Fig. S2; Supplementary Table S3). The majority of these cells were positive for *Spink4*, but did not produce the other typical secretory cell markers *Atoh1* and *Spdef*, which is why we named this cluster Atoh1^-^/Spdef^-^ cells. Interestingly, scRNA velocity analysis revealed the origin of the cells in cluster 14. Another notable observation was that the production of Tcf/Lef factors by the cells in clusters 9 and 14 was essentially undetectable (Fig. 2B). This could indicate that these cells are Tcf4 independent and either remain in the epithelium after *Tcf7l2* cKO or are newly generated in the regenerating epithelium, which would explain their higher abundance in the *Tcf7l2^flox/flox^* sample (Fig. 2A). The cells in cluster 15 mainly produced immune cell markers. These cells (according to the expression profiles CD3/CD8^+^ T cells; Supplementary Table S3) were possibly (co-)isolated with intestinal cells.

**Figure 2.**
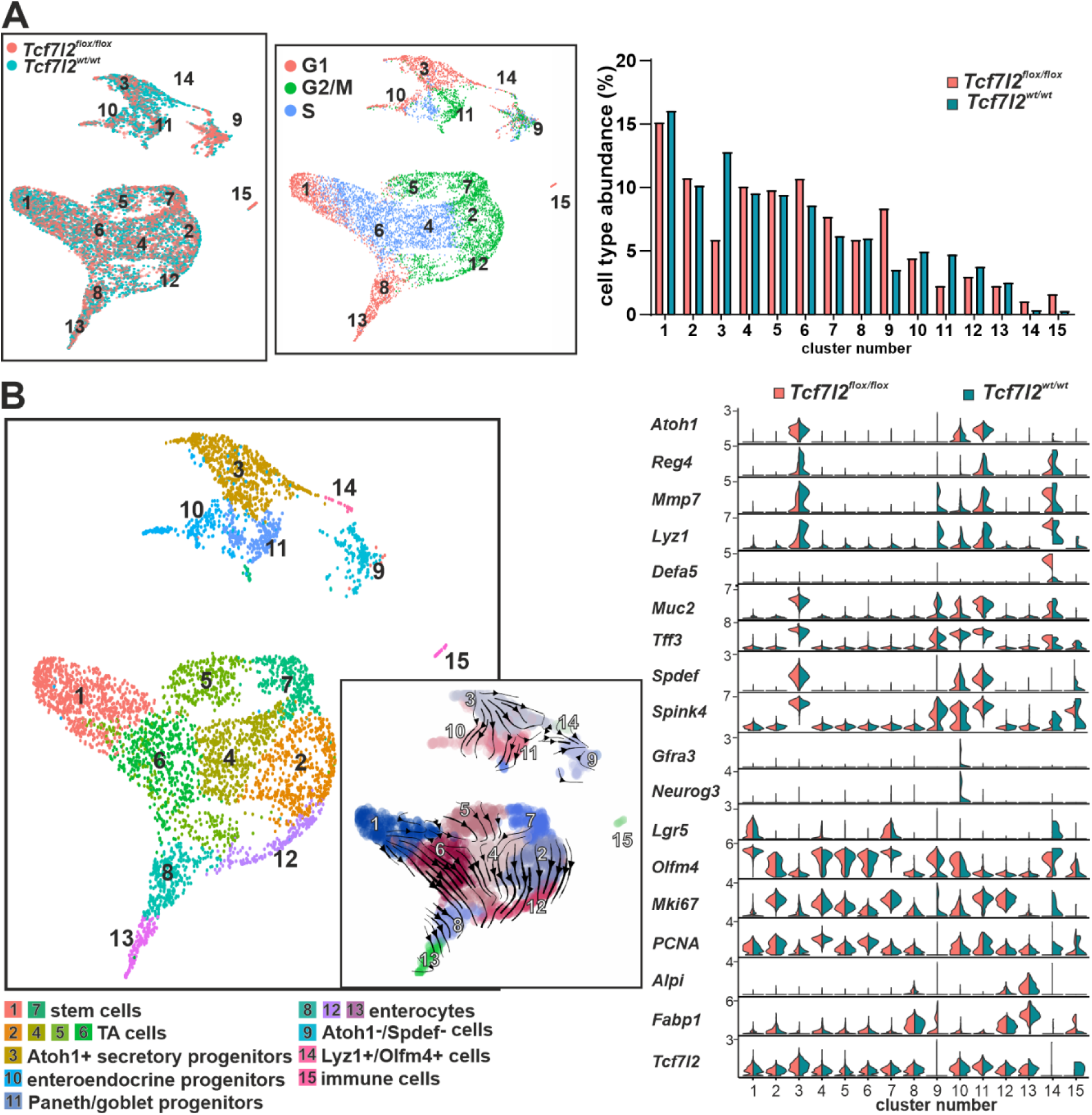
Transcriptome profiling of proliferating intestinal epithelial cells reveals cell type-specific gene expression and lineage dynamics in *Tcf7l2* genotype variants. A) Single-cell transcriptome analysis of intestinal epithelial cells to compare gene expression in *Tcf7l2^wt/wt^/Mki67^RFP/RFP^/Villin-CreER^T2^*(*Tcf7l2^wt/wt^*) and *Tcf7l2^flox/flox^/Mki67^RFP/RFP^/Villin-CreER^T2^*(*Tcf7l2^flox/flox^*) mice. The animals were administered tamoxifen and after 7 days, proliferating (i.e., Mki67-RFP-positive) epithelial cells were isolated and analyzed. The diagrams on the left show a UMAP (Uniform Manifold Approximation and Projection) plot colored according to genotype or cell cycle phase. The bar chart on the right shows the frequency of cell types in the clusters of the two scRNA-seq samples. The percentages are plotted so that the total number of cells in each sample is 100%. B) The single-cell transcriptome analysis shows different epithelial cell types of the intestine. The left diagram shows a UMAP visualization in which the cells are color-coded according to their identified cell type. The inset diagram is an overlay of the UMAP clusters with arrows representing the lineage relationships between the cell types. The violin diagram on the right shows the differential expression of key lineage markers in the identified clusters compared between *Tcf7l2^wt/wt^*and *Tcf7l2^flox/flox^* mice; the level of gene expression is indicated on the y-axis. Alpi, alkaline phosphatase, intestinal; Atoh1, atonal bHLH transcription factor 1; Defa5, defensin alpha 5; Fabp1, fatty acid binding protein 1; Gfra3, GDNF family receptor alpha 3; Lgr5, leucine-rich repeat-containing G protein-coupled receptor 5; Mmp7, matrix metallopeptidase 7; Mki67, proliferation marker Ki-67; Muc2, mucin 2; Neurog3, neurogenin 3; Olfm4, olfactomedin 4; Reg4, regenerating family member 4; Spdef, SAM pointed domain containing ETS transcription factor; Spink4, serine peptidase inhibitor Kazal type 4; Tff3, trefoil factor 3.

### Control of secretory cell type switching between Paneth and goblet cells by the *Tcf7l2* gene in Defa6^+^ cells of the small intestine

In our model of intestinal crypt regeneration after partial loss of Tcf4, we observed misplaced cells that were positive for the Paneth cell marker lysozyme. This finding underlines the importance of the Wnt signaling pathway not only for homeostatic self-renewal but also for Paneth cell maturation, as van Es and colleagues have previously noted (37). Thus, we investigated the effects of *Tcf7l2* gene inactivation using the *Defa6-iCre* mouse strain developed to target Paneth cells in the intestine (38). In *Rosa26-tdTomato*/*Defa6-iCre* mice, we observed co-localization of tdTomato-labeled (Defa6-tdTom) cells with a Paneth cell marker lysozyme at the base of small intestinal crypts (Fig. 3A). Homozygous knockout of *Tcf7l2* in Defa6^+^ cells resulted in the absence of lysozyme granules and morphological changes indicative of vacuolization. In addition, the number of lysozyme-positive cells decreased significantly throughout the small intestine, especially in the ileum (Fig. 3B). Although the overall architecture of the intestinal epithelium remained intact, a slight increase in proliferative activity was observed in the crypts. In addition, the area labeled with Olfm4, which denotes the stem cell compartment (27), increased in 2-3 months old mice, while it decreased in 1-year old mice (Supplementary Fig. S3A), indicating a possible decrease in stem cell reserves due to the "efforts" to compensate for the loss of Paneth cells and maintain epithelial homeostasis (39). Tracking of Defa6^+^ cells by tdTomato fluorescence showed that inactivation of *Tcf7l2* resulted in migration of Defa6-tdTom cells from the crypts into the villi, similar to the migration observed in other differentiated intestinal epithelial cells. These cells did not exhibit the structure characteristic of Paneth cells; in addition, they produced the goblet cell marker Muc2 (see Fig. 3C).

**Figure 3.**
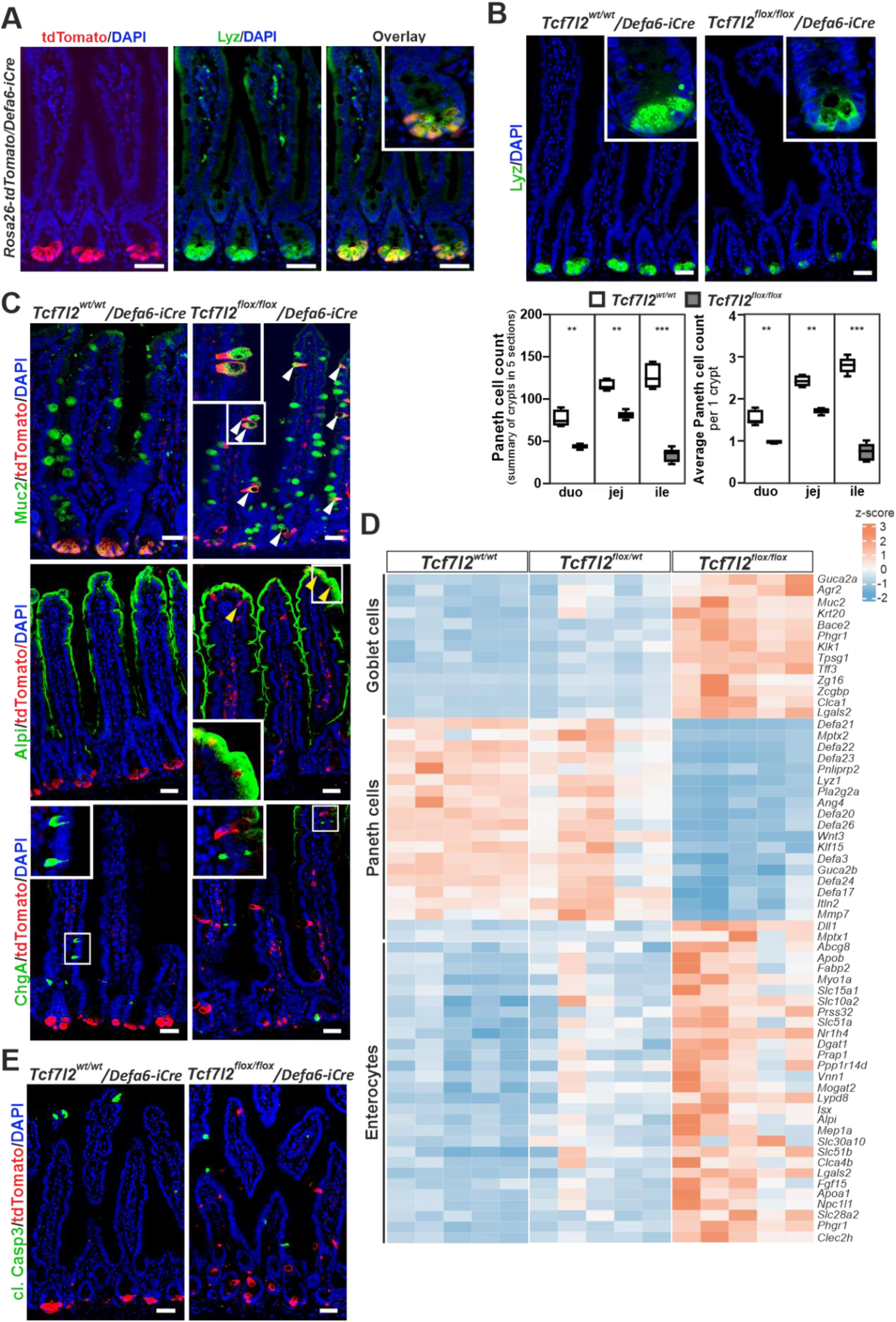
Comparative analysis of the morphology and distribution of Paneth cells in *Tcf7l2* wild-type and knockout mice. Mouse lines *Tcf7l2^wt/wt^/Defa6-iCre* and *Tcf7l2^flox/flox^/Defa6-iCre* were used to analyze Defa6-tdTom cells after crossing with *ROSA26-tdTomato* reporter mice. A) Confocal microscopy images show the co-localization of tdTomato (red signal) with lysozyme (Lyz; green signal) in intestinal sections. The overlay highlights the extent of co-localization. Scale bar: 50 µm. Magnified images are shown in the insets. B) The upper panel shows Lyz expression in Paneth cells. The insets show the detailed cellular morphology and highlight the differences between Tcf4 wt (*Tcf7l2^wt/wt^*) and knockout (*Tcf7l2^flox/flox^*) conditions. The lower panels show quantitative analyses of the number of Paneth cells (based on Lyz positivity) in the duodenum (duo), jejunum (jej) and ileum (ile). Statistical significance was determined using a one-way ANOVA test; **p < 0.01; ***p < 0.001. C) Top images show mislocalized Muc2-positive *Tcf7l2^flox/flox^*/Defa6-tdTom cells (indicated by white arrowheads); middle: presence of Alpi in a small subset of *Tcf7l2^flox/flox^/Defa6-tdTom* cells on the villi (indicated by yellow arrowheads), which lack the morphology of Paneth cells. Bottom: lack of chromogranin A (ChgA) staining in Defa6-tdTom cells under both conditions. Scale bar: 50 µm. D) Heatmap of differentially expressed genes obtained by bulk RNA-seq from Defa6-tdTom cells with different genetic contexts (as indicated). Each row represents a gene and each column represents a specific cell type. Color intensity (blue to red) indicates the Z-score of gene expression levels and highlights differences and patterns of up- or downregulation. A complete list of differentially expressed genes can be found in Supplementary Table S4. E) Staining of cleaved caspase 3 (cl. Casp3) indicates apoptotic activity at the apical tips of small intestinal villi. Scale bar: 50 µm.

Early signs of this cell migration were observed in the postnatal intestine as early as 15 days after birth (Supplementary Fig. S3B), which is consistent with the developmental timing of Paneth cells in the mouse small intestine (40). Cells located in the lower part of the small intestinal crypts, including Paneth cells, produce surface marker CD24 (41). Interestingly, FACS analysis showed virtually no CD24-negative Defa6-tdTom cells in wt mice (genotype: *Tcf7l2^wt/wt^/Rosa26-tdTomato/Defa6-iCre*). In contrast, CD24-negative Defa6-tdTom cells were observed in *Tcf7l2^flox/flox^/Rosa26-tdTomato/Defa6-iCre* mice (Supplementary Fig. S4A). Isolation and genomic DNA analysis of both CD24-positive and CD24-negative tdTomato^+^ populations confirmed the presence of the recombinant *Tcf7l2^del(Ex5)^* allele in *Tcf7l2* cKO mice, with increased presence in CD24-negative cells, indicating ongoing recombination and a decrease in functional Tcf4 protein in maturing Paneth cells (Supplementary. Fig. S4B). Quantitative RT-PCR analysis showed that the expression of *Tcf/Lef* genes was highest in wt mice in CD24-positive Defa6-tdTom cells, presumably Paneth cells. We detected a gradual decrease in *Tcf7l2* mRNA in both CD24-positive and CD24-negative Defa6-tdTom cell populations in *Tcf7l2* cKO mice. Interestingly, CD24^+^ Defa6-tdTom cells exhibited a significant decrease in expression of the *Tcf7* gene (encoding Tcf1), whereas expression of the paralogous *Tcf7l1* gene (encoding Tcf3) was increased. However, the decreased expression of the Wnt-responsive genes *Axin2* and naked cuticle homolog 1 (*Nkd1*) suggests that other members of the Tcf/Lef family are unable to maintain physiologic levels of Wnt signaling in the absence of functional Tcf4. Consequently, CD24-negative Defa6-tdTom cells also lost expression of crypt base-specific stem cell genes such as *Lgr5*, tumor necrosis factor receptor superfamily, member 19 (*Tnfrsf19*) (42), and *Olfm4* as well as the Paneth cell-specific genes *Defa24* and *Lyz1*. In contrast, we observed a strong increase in the expression of mRNAs characteristic of goblet cells, such as chloride channel accessory 1 (*Clca1*), *Muc2* and *Tff3* (43). In addition, the expression of genes encoding enterocyte markers such as *Alpi*, *Fabp1* and *Sis* was increased (35, 36) (Supplementary Fig. S4C). Accordingly, histologic staining for the major intestinal cell type markers revealed a small number of Defa6-tdTom cells at the tips of some villi in *Tcf7l2^flox/flox^/Rosa26-tdTomato/Defa6-iCre* mice. These cells lacked the vacuolar pattern typical of goblet cells and (some) showed strong staining for Alpi. In contrast, staining for chromogranin A (ChgA), which is specific for enteroendocrine cells, never overlapped with tdTomato labeling (Fig. 3C). To confirm these results, we performed bulk RNA sequencing on Defa6-tdTom cells from combinations of *Tcf7l2* alleles, i.e., homozygous wt, heterozygous and homozygous cKO. While cells with heterozygous loss of *Tcf7l2* were almost identical in expression to wt cells, homozygous deletion of *Tcf7l2* resulted in marked changes in gene expression (Supplementary Table S4). According to the Panglao database of cell type-specific gene expression (35), this was characterized by a marked loss of genes specific to Paneth cells and upregulation of genes characteristic of goblet cells and enterocytes (Fig. 3D). In addition, visualization of cleaved caspase 3 showed that apoptosis in Defa6-tdTom cells occurs mainly at the villus tips regardless of Tcf4 status, indicating normal homeostatic cell renewal (Fig. 3E).

To comprehensively characterize the development of the Paneth cell lineage in the intestine and to show to what extent the loss of Tcf4 leads to its alteration, we performed scRNA-seq on Defa6-tdTom cells from the small intestine of *Tcf7l2^wt/wt^/Rosa26-tdTomato/Defa6-iCre* (Tcf4 wt) or *Tcf7l2^flox/flox^/Rosa26-tdTomato/Defa6-iCre* (Tcf4 cKO) mice. Since we observed a low cell yield in samples with the Tcf4 cKO genotype, we isolated Defa6-dTom cells of both the Tcf4 wt and Tcf4 cKO genotypes together. We then compared the results obtained with this “mixed” sample (Sample 2 in Fig. 4A) with the results obtained when only Defa6-tdTom Tcf4 wt cells were isolated and analyzed (Sample 1 on Fig. 4A). The combined analysis of both datasets showed that Lgr5/Olfm4-positive proliferating (PCNA- and Mki67-positive) cells or rapidly dividing cells (producing large amounts of ribosomal proteins) (44, 45) were present in three clusters (6, 8 and 3, respectively). The cells in these clusters expressed target genes of the canonical Wnt signaling pathway *Axin2* and *Sp5* (46), indicating active Wnt signaling (Fig. 4B). Since *Lyz1* (and *Spink4*) was produced at significant levels in almost all cells in the scRNA-seq analysis, we were hesitant to name these cells as stem/TA cells. Therefore, we named clusters 3, 6 and 8 as "Lgr5^+^/Olfm4^+^ cells". It should be noted that the transcriptional regulator achaete-scute family bHLH transcription factor 2 (*Ascl2*) of intestinal stem cells (47) was preferentially produced in the cells of clusters 6 and 8 (Supplementary Fig. S5AB). The cells in the other eight clusters were predominantly in the G1 phase of the cell cycle. Cluster 10 included enterocytes [expression: angiotensin-converting enzyme 2 (*Ace2*), *Alpi*, *Fabp1/2*, keratin 20 (*Krt20*), and *Sis*]. Paneth cell progenitors expressing *Atoh1* and transcription factor *Sox9*, which is critical for Paneth cell differentiation (48), and general Paneth cell marker genes such as *Mmp7* and *Lyz1*, and defensins were included in cluster 1. The clusters with significantly reduced cell numbers in the combined sample (Tcf4 wt/cKO) were 4 and 5. These cell clusters represent mature Paneth cells expressing a number of defensins, *Lyz1* and mucosal pentraxin 2 (*Mptx2*) (Fig. 4ABC). In addition, three clusters (2, 7 and 11) represented the goblet cell lineage. Cluster number 7 represented mature goblet cells that were positive for a number of goblet cell markers [*Clca1*, *Muc2*, Fc-γ binding protein (*Fcgbp*), *Spdef*, *Tff3*] and also for molecules that indicate the position of the cell on the villi, in particular *Krt20*, zymogen granule protein 16 (*Zg16*), and indoleamine 2,3-dioxygenase 1 (*Ido1*) (35, 49). This cluster, together with clusters 3 and 6 (actively dividing cells), was enriched in the sample with the Tcf4 wt/cKO cell mixture (Fig. 4B). Cluster 2 probably contained goblet cell progenitors, which, in addition to the markers for these cells, also produce the *CD24a* gene (encoding the surface marker CD24), which, as already mentioned, is typically produced in the lower part of the crypts. The least abundant cluster was cluster 11, which was characterized by the absence of transcripts encoding members of the Tcf/Lef family genes (Fig. 4B). The cells in this cluster, named “Tcf7l2^-^ secretory progenitors”, dominantly expressed markers for goblet cells as well as *Krt19*, which marks the crypt compartment of TA cells (50). In addition, the cells in this cluster produced fewer transcripts encoding the transcription factors *Atoh1* and *Spdef*. Cluster number 9 contained transcripts encoding intestinal stem cell marker leucine-rich repeats and immunoglobulin-like domains 1 (*Lrig1*) and *Lgr5* homolog *Lgr4*. However, other genes expressed by the cells in this cluster did not indicate that they were intestinal stem cells (Fig. 4C). The cells in cluster 9 differed from those in the other clusters by the expression of gene *Sntb1* (Fig. 4C; Supplementary Table S5), which codes for endocytic adaptor syntrophin beta 1 (51). Therefore, we named this cluster "Sntb1^+^ cells". Interestingly, we noted high expression of epidermal growth factor (*Egf*) in this cluster. As goblet cell differentiation is dependent of mitogen activated protein kinase (MAPK) pathway (52) and we noted Egf receptor expression preferentially in the goblet cell linage (Supplementary Table S5), these cells could represent a subpopulation of cells involved in Paneth/goblet cell fate determination.

**Figure 4.**
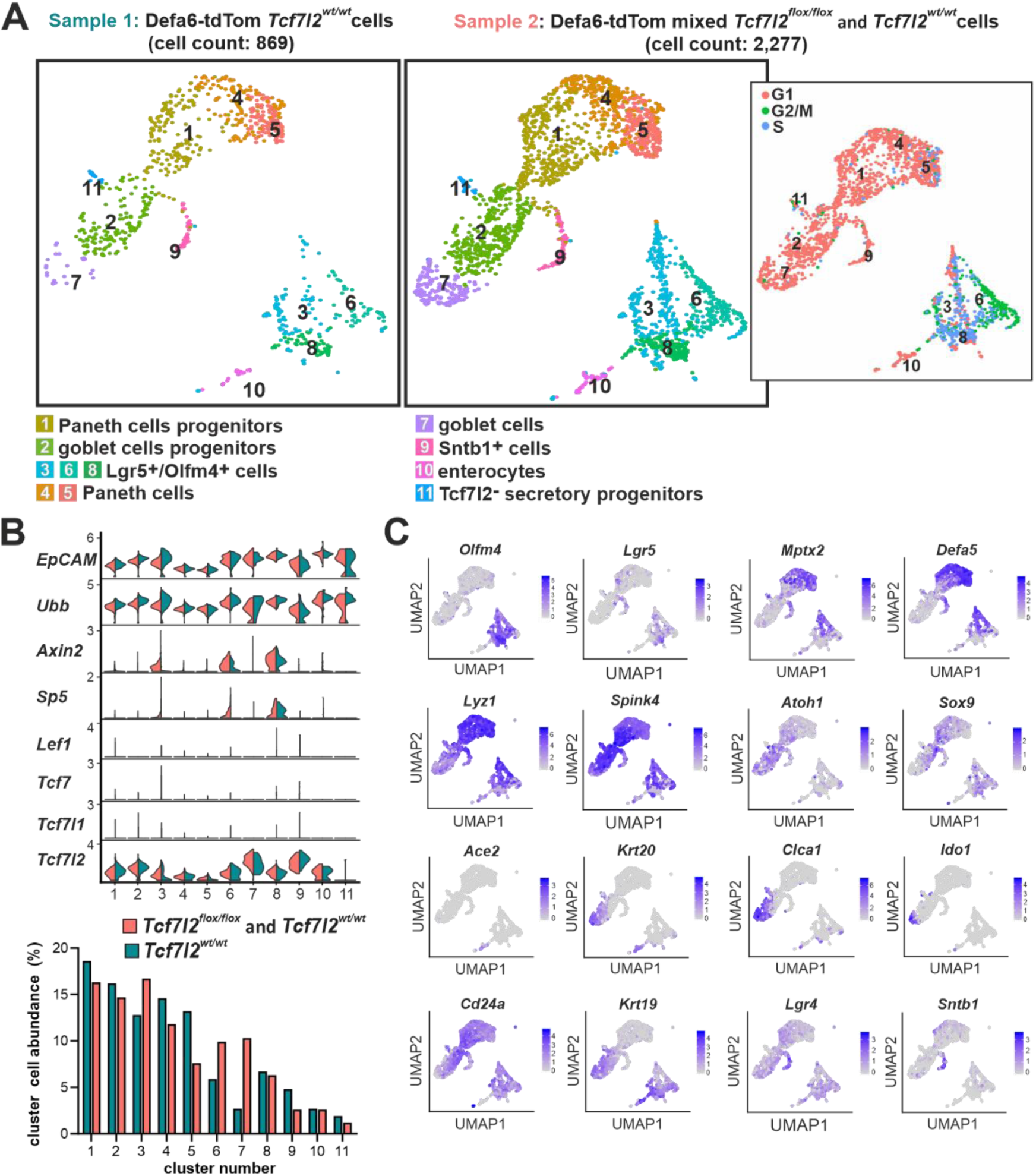
Cells after loss of Tcf4 leave the Paneth cell lineage and acquire gene expression characteristics of goblet cells and enterocytes. (A) Single-cell RNA-seq data of Defa6-tdTom cells from the small intestine of different Tcf7l2 genotypes. Left, UMAP visualization of scRNA-seq clustering of Defa6-tdTom cells from the small intestine of mice with the wt *Tcf7l2* gene (*Tcf7l2^wt/wt^*, sample 1) or Defa6-tdTom cells in a mixed sample of *Tcf7l2^wt/wt^* and *Tcf7l2^flox/flox^*cells (sample 2; right). Cell cycle analysis of Defa6-tdTom cells combined from both samples is shown in the inset. The cell clusters are numbered according to the number of cells. Cell types were assigned based on the genes specifically expressed in each cluster (see Supplementary Table S5 for a complete list of genes). (B) Top, violin plots showing the expression level of *EpCAM* and *Ubb* genes as well as genes of the Tcf/Lef family in both Defa6-tdTom scRNA-seq samples. The expression levels of *Axin2* and *Sp5*, which indicate the status of Wnt signaling, are also shown in the identified cell clusters from panel A. Bottom, frequency of cell types in the clusters of the two scRNA-seq samples. The percentages are plotted so that the total number of cells in a respective sample is 100%. Lef1, lymphoid enhancer-binding factor 1. (C) UMAP plots showing the expression patterns of the different marker genes. The plots were derived from both datasets. Ace2, angiotensin-converting enzyme 2; Clca1, chloride channel accessory 1; Ido1, indoleamine 2,3-dioxygenase 1; Krt19/20, keratin 19/20; Mptx2, mucosal pentraxin 2; Sntb1, syntrophin beta 1; Sox9, sex-determining region Y (SRY)-box 9.

In addition, distribution of differentially expressed genes (determined by bulk RNA-seq) in the individual cell clusters identified in the scRNA-seq analysis confirmed lower abundance of the Paneth cell lineage and increased presence of the goblet cell lineage observed in the Tcf4 wt/cKO compared to the Tcf4 wt sample (Supplementary Fig. S5C). Among the most altered biological processes in gene ontology (GO), we found signaling pathways related to immune and antimicrobial responses. In addition, pathways related to the localization and targeting of proteins in the endoplasmic reticulum (ER) or endoplasmic reticulum-associated degradation (ERAD) (53) were also significantly affected, indicating the presence of actively secreting cells in the samples studied (Supplementary Fig. S5D).

### Multiple source cells of the Wnt3 ligand in the epithelium of the small intestine

Paneth cells not only provide antimicrobial protection, but also create a microenvironment known as a niche for intestinal stem cells and produce Wnt3, which is important for maintaining the "stemness" of ISCs (41). The competing activities of the Wnt and Notch signaling cascades are required for ISC maintenance and determine the cell fate during differentiation. While induction of Notch signaling drives differentiation into the absorptive lineage, the absence of Notch signaling causes progenitor cells to differentiate into the secretory lineage (54). An interesting, if somewhat expected fact was that the secretory progenitor cells, regardless of the type of compartment, expressed delta-like canonical Notch ligand (*Dll*) *1/4* of the Notch signaling pathway. In contrast, cells in the stem cell compartment produced the receptor of this pathway, *Notch2*, and, according to the expression of the target gene of this pathway hes family bHLH transcription factor 1 (*Hes1*), actively signaled from the receptor (37). A similar situation was observed for Wnt signaling. The *Wnt3* ligand was expressed in secretory progenitor cells, while the target genes of this pathway *Axin2*, *Tnfrsf19* and ring finger protein 43 (*Rnf43*) (55) were mainly produced in stem cells and to some extent also in the progenitor compartment (Fig. 5A). We also detected *Wnt3* expression in most Defa6-tdTom cell types, i.e. in both secretory progenitor cells and Paneth cells. In addition to *Wnt3*, we observed expression of the canonical ligand *Wnt7b* (56) and the non-canonical *Wnt11*, (57) (58). Bulk RNA-seq showed that inactivation of the T*cf7l2* gene in Defa6-tdTom cells resulted in decreased expression of these Wnts (Fig. 5B and Supplementary Table S4).

**Figure 5.**
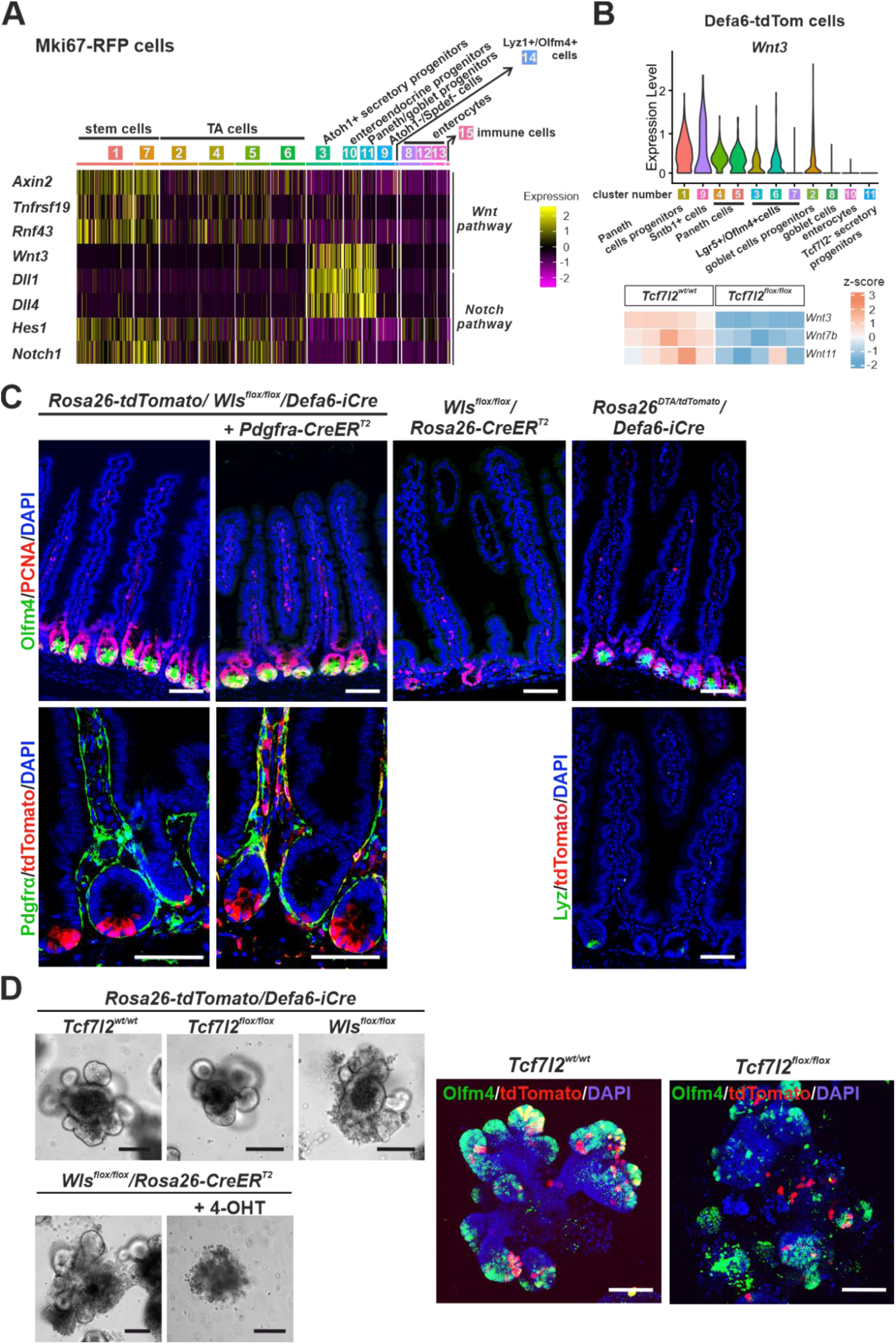
Different cell types that produce Wnt ligands in the small intestinal epithelium. A) Heatmap with scaled expression of genes involved in the Wnt (top) and Notch (bottom) signaling pathways. Expression levels are color-coded, with yellow indicating high expression and purple indicating low expression. Dll1/4, Delta-like canonical Notch ligand 1/4; Hes1, Hes family basic helix-loop-helix (bHLH) transcription factor 1; Notch1, Notch homolog protein 1; Rnf43, ring finger protein 43; Tnfrsf19, tumor necrosis factor receptor superfamily, member 19. B) Top, violin plot showing the expression levels of Wnt3 in different cell clusters of Defa6-tdTom cells. Bottom, heatmap showing the expression of indicated Wnt ligands in Tcf4 WT Defa-tdTom cells. In cells with homozygous Tcf4 cKO, downregulation of all Wnt ligands was detected. C) Immunohistochemical analysis of the crypt compartment after genetic knockout of the *Wls* gene, 8 days after recombination, using indicated Cre recombinase-expressing mouse strains. Left and center, the images showing the absence of Olfm4 and reduced PCNA staining observed only after inhibition of Wnt ligand secretion in all epithelial and subepithelial cells. Right, additional panel of Muc2 and tdTomato staining highlighting the mucus-producing goblet cells in the intestinal crypts and showing the effects of Paneth cells depletion using diphtheria toxin A (DTA). Bottom, documentation of recombination in Defa6^+^ and Pdgfrα^+^ cells using tdTomato reporter protein fluorescence. The scale bars correspond to 0.15 mm. Wls, Wntless. D) Left, growth of intestinal organoids from crypts containing *Tcf7l2* wt, *Tcf7l2* cKO, or *Wls* cKO in Defa6-tdTom cells (upper panel) or *Wls* cKO in all cells induced by administration of 4-hydroxytamoxifen (4-OHT) in culture medium (bottom panel). Right, representative fluorescence images of organoids showing Olfm4-positive crypt compartments and Defa6-tdTom cells. The scale bars correspond to 0.1 mm.

The loss of Paneth cells after inactivation of the *Tcf7l2* gene had no effect on epithelial renewal, which was expected since it has already been predicted and subsequently demonstrated that Paneth cells are not the only source of Wnt ligands in the intestine (59–61), see further. To achieve complete loss of Wnt ligand secretion in Defa6-tdTom cells, we used a mouse strain that allows conditional inactivation of the Wntless (*Wls*) gene, which encodes a transmembrane protein essential for Wnt ligand secretion from the cell (62). Similar to the loss of Wnt ligand expression in Tcf4-deficient Paneth cells, *Wls^flox/flox^* in Defa6^+^ cells had no effect on the number of intestinal stem cells or homeostatic renewal of the epithelium. Similarly, no morphological changes of the epithelium were observed after depletion of Paneth cells in *Rosa26^DTA/tdTomato^/Defa6-iCre* mice, which allow production of diphtheria toxin A (DTA), a so-called suicide gene, from the *Rosa26* locus (63) (Fig. 5C). Recently, subepithelial mesenchymal cells have been shown to serve as a secondary source of Wnt signaling for intestinal stem cells (61, 64). Therefore, we subsequently inhibited Wnt ligand secretion in mesenchymal cells using mice that produce CreER^T2^ recombinase in mesenchymal cells producing platelet-derived growth factor receptor alpha (*Pdgfrα*) (65, 66). Surprisingly, the simultaneous blocking of Wnt ligand secretion in Defa6^+^ and Pdgfrα^+^ cells did not lead to a loss of intestinal stem and proliferating cells. In contrast, inactivation of the *Wls* gene with the *Rosa26-CreER^T2^* driver, which enables deletion of the floxed allele in all cell types, led to a significant reduction in the number of proliferating cells in the crypts of the small intestine (Fig. 5C).

We then established organoid cultures and compared the effects of inactivating the *Tcf7l2* gene in Defa6-tdTom cells, either with the *Defa6-iCre* transgene or with inactivation of the *Wls* gene with the *Defa6-iCre* or *Rosa26-CreER^T2^* driver. While organoids with inhibited Wnt secretion in all cells collapsed after the first passage, organoids with inactivated *Tcf7l2* or *Wls* in Defa6^+^ cells grew and showed a branched morphology similar to wt organoids for at least five passages. However, the "crypts" of organoids with the *Tcf7l2* cKO allele were partially disorganized and had fewer Olfm4-positive cells. In addition, tdTomato-positive cells were more centrally located, i.e., in the organoid region that contained more derived cell types (Fig. 5D). Thus, this phenotype resembled the situation observed in *in vivo* experiments.

### A gene expression program for the production of antimicrobial peptides is initiated in early colon adenomas

We observed that Tcf4 deficiency led to decreased expression of antimicrobial peptides in Defa6^+^ cells of the small intestine. This prompted us to ask whether, conversely, the expression of antimicrobial peptides in tumors could be increased due to abnormally enhanced Wnt signaling. Following *Apc* ablation in the colon, there was a remarkable increase in cell proliferation, colonic crypt expansion, and expression of Tcf4 (Supplementary Fig. S6A). In addition, during the initial phase of adenoma formation in the colon, the first significant response of the hyperplastic colonic epithelium was the synthesis of a plethora of antimicrobial peptides, including lysozyme and a variety of α-defensins, which could be detected as early as two days after tamoxifen-induced *Apc* gene inactivation (19) (Fig. 6A). Key biological processes in the tissue at this time also included innate immune responses and the defense mechanism against bacteria. On the fourth day after *Apc* loss, these responses were overshadowed by transcriptional shifts and DNA replication processes, indicating increased proliferation of the epithelium (Supplementary Figure S6B).

**Figure 6.**
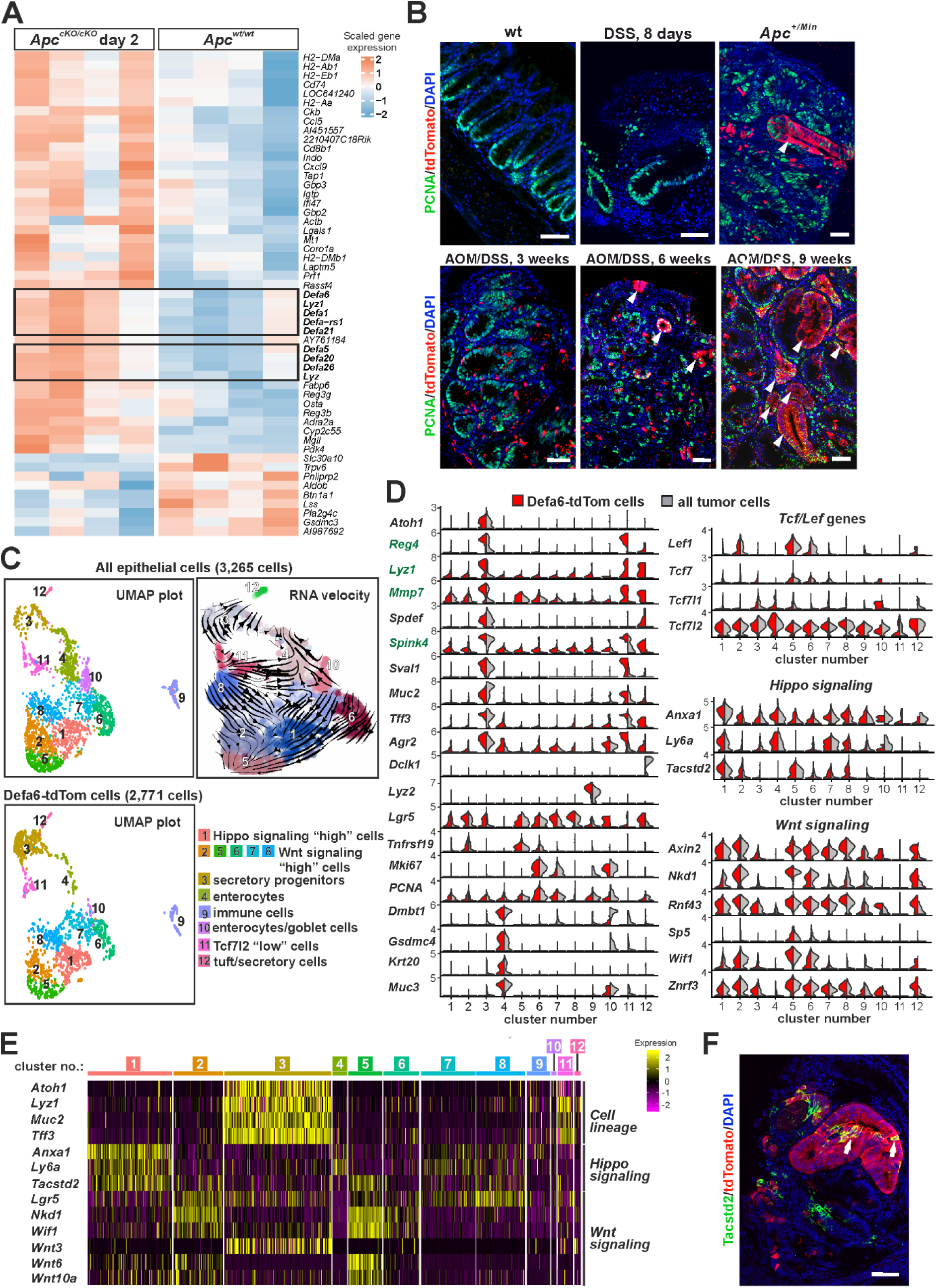
Activation of α-defensin expression during the development of colonic adenomas. A) Microarray expression profiling of colonic crypts. The heatmap shows differentially expressed genes with color scaling indicating the expression level in tissue 2 days after cKO of the adenomatous polyposis coli (*Apc*) gene compared to tissue with the wild-type (wt) *Apc*. Significant genes (adjusted p-value < 0.05 and |log_2_ fold change (FC) > 1) are shown. Paneth cells-specific genes are boxed. See Supplementary Fig. S5B for enriched Gene Ontology (GO) biological processes. B) The appearance of Defa6-tdTom cells is associated with tumorigenesis in the colon. Representative fluorescence microscopy images of cells stained with an antibody against PCNA (green signal) and tdTomato (red signal). Note that the red fluorescent signal was observed either in mice treated with the mutagen azozymethane (AOM) in combination with colitis-inducing sodium dextran sulfate (DSS) or in the colon tumor that developed in *Apc^+/Min^* mice. In contrast, "red" cells are present neither in the healthy colonic epithelium (wt) nor in the epithelium 8 days after induction of colitis by DSS treatment, i.e., without mutagen (pre-)treatment (top right image). Notice the red glandular structures that are formed 6 and 9 weeks after AOM/DSS treatment and in tumors of the *Apc^+/Min^* mice (white arrowheads in the upper right and bottom middle and right image). Samples were counterstained with 4′,6-diamidino-2-phenylindole dihydrochloride (DAPI; blue nuclear signal). Scale bar: 150 µm. C) Single-cell RNA sequencing (scRNA-seq) analysis of epithelial cells from dissected colon tumors in *Apc^+/Min^* mice. Cell clusters are numbered according to cell abundance; UMAP visualizations show all epithelial tumor cells (left) and Defa6-tdTom cells (right). The middle diagram is an overlay of the UMAP clusters with arrows representing the lineage relationships between the cell types (RNA velocity). For better visibility, the diagram has been expanded spatially and cluster 9 (immune cells) is not shown. A complete list of cluster marker genes can be found in Supplementary Table S6. D) Left, the expression of marker genes in different cell clusters is shown in the violin plots, with the level of gene expression indicated on the y-axis. Genes with increased expression in Defa6-tdTom tumor cells are shown in green. Right, analysis of the components of the Wnt signaling pathway in individual cell clusters. Each violin plot represents the mean expression level for the Wnt signaling target genes, or the genes encoding the nuclear mediators of the Wnt signaling pathway Tcf/Lef or genes activated upon Hippo pathway inhibition. Expression in all epithelial tumor cells is shown in gray, Defa6-tdTom tumor cells are shown in red. Agr2, anterior gradient 2; Anxa1, annexin A1; Dclk1, doublecortin-like kinase 1; Dmbt1, deleted in malignant brain tumors 1; Gsdmc, gasdermin C3; Ly6a, lymphocyte antigen-6; Nkd1, naked cuticle homolog 1; Sval1, seminal vesicle antigen-like 1; Tacstd2; tumor-associated calcium signal transducer 2; Wif1, Wnt inhibitory factor 1; Znrf3, zinc and ring finger 3. E) Heatmap showing scaled expression of indicated genes in sorted Defa6-tdTom tumor cells; cluster numbers were obtained from the combined analysis of total tumor cells and Defa6-iCre-labeled cells. Expression levels are color-coded, with yellow indicating high expression and purple indicating low expression. F) Representative fluorescence microscopy images of cells stained with an antibody against Tacst2 (green signal) and tdTomato (red signal). Note the Tacstd2-positive cells in the red glandular structure (white arrows). Scale bar: 150 µm.

Given the reported expression of the α-defensin-6 gene (*DEFA6*) in human adenomas (67) and the potential role of the antimicrobial response in the development of colorectal tumors, we wanted to investigate this occurrence further using *Rosa26-tdTomato/Defa6-iCre* mice. In adult mice, red fluorescence was not observed in either healthy colonic epithelium or epithelium 8 days after induction of colitis by DSS treatment. Therefore, activation of the α-defensin program is involved neither in the homeostatic turnover of the intestinal epithelium nor in the regeneration of the crypt compartment after injury. In contrast, we detected the presence of Defa6-tdTom cells in mice when DSS administration was preceded by administration of mutagen AOM and in colon adenomas of mice developing multiple intestinal neoplasia (Min). The *Apc^+/Min^* mice carry a single nucleotide mutation in one allele of the *Apc* gene, which leads to the production of a non-functional truncated protein. As a result of the random inactivation of the wt *Apc* allele, the mice develop numerous adenomas in adulthood, predominantly in the small intestine and in some individuals also in the large intestine (68). Interestingly, the adenomas examined three weeks after AOM/DSS treatment showed only scattered Defa6-tdTom cells. In contrast, the more advanced colon adenomas of *Apc^+/Min^* mice showed clusters or glands of Defa6-tdTom cells in addition to scattered cells. However, six weeks after AOM/DSS treatment, glandular structures consisting of Defa6-tdTom cells began to form (Fig. 6B).

We used FACS to isolate either all epithelial cells (EpCAM^+^) or Defa6-tdTom/EpCAM^+^ cells from dissected colon tumors; the sorting strategy is shown in Supplementary Fig. S7A. We then analyzed these cells by scRNA-seq. The obtained datasets were merged and clusters were identified based on the previously described cell type-specific gene expression profiles (69–71) (Fig. 6CD). A complete list of genes expressed in the individual clusters can be found in Supplementary Table S6.

As expected, many of the profiled cells were proliferating, as shown by the expression of *PCNA* and/or *Mki67*, albeit to varying degrees (Fig. 6D). Almost all cell clusters expressed the transcription factor *Sox9*, which has previously been associated with tumorigenesis in the colon epithelium (72, 73) (not shown). Another gene that was expressed almost in all clusters was *Lgr5*. In several clusters (nos. 2, 5 and 6), we observed co-expression of *Lgr5* and another intestinal stem cell marker, *Tnfrsf19*. The molecular basis of tumorigenesis in *Apc^+/Min^* mice is the abnormal activation of the canonical Wnt signaling pathway mediated by β-catenin (reviewed in (74)). Logically, one would expect tumor cells to produce more Wnt signaling target genes. This assumption was supported by the observation that the majority of cells in the clusters examined expressed the Wnt signaling target gene *Axin2*, *Rnf43*, *Sp5*, Wnt inhibitory factor 1 (*Wif1*), and zinc and ring finger 3 (*Znrf3*), which had previously been identified as a target gene in intestinal tumors (75) (Fig. 6DE). In clusters 2, 5-8, the relatively highest amounts of the canonical target gene *Axin2* and of the *Rnf43* gene were produced. Apart from the overproduction of *Lgr5*, these clusters did not show differential expression of the so-called cell line-specific genes, which is why we designated these clusters as Wnt signaling "high" cells. Interestingly, RNA velocity analysis revealed a cellular trajectory of cells in the above clusters toward cluster 5, and cells in this cluster showed the highest expression of many target genes of the Wnt signaling pathway, including *Lef1*, which is also activated by the Wnt signaling pathway in intestinal epithelial cells (76). On the way to cluster 5, there were also cells from cluster 1, which was characterized by an increased expression of genes regulated by the Hippo signaling pathway. These included the annexin A1 gene (*Anxa1*) and genes coding for the so-called oncofetal (regenerative) stem cell antigen Sca1 (encoded by lymphocyte antigen-6; *Ly6a*) and tumor-associated calcium signal transducer 2 (*Tacstd2*) (Fig. 6DE) (77–79). Interestingly, the cells positive for Tacstd2 formed clusters in the tumor including glandular structures labeled with *Defa6-iCre*; however, the scattered Defa6-tdTom cells were predominantly Tacstd2-negative (Fig. 6F). Furthermore, our expression analyses of all four members of the Tcf/Lef family members confirmed previous findings (see (80)) that Tcf4 is the predominant nuclear mediator of the Wnt signaling pathway in the colon (Fig. 6D). Cell clusters that were the most overrepresented in samples containing all epithelial tumor cells were clusters 4 and 10, with cells in cluster 4 expressing *Krt20*, *Muc3*, and enterocytes markers deleted in malignant brain tumors 1 (*Dmbt1*) and gasdermin C3 (*Gsdmc*). Gene set enrichment analysis using the Enricher Web platform (81, 82) showed that this cluster indeed contained enterocytes. This cluster, although less abundant, was also clearly defined in the Defa6-tdTom cells, and RNA velocity analysis showed it to be terminal. Cluster 10 appeared more heterogeneous and contained markers of enterocytes and goblet cells. Another heterogeneous cluster was cluster 11, which was characterized by the expression of secretory lineage marker *Spink4* and production of markers for Paneth cells (*Lyz1*, *Mmp7*) and goblet cells [*Agr2*, *Muc2*, seminal vesicle antigen-like 1 (*Sval1*) and *Tff3*]. The common feature of the cells in this cluster was the absence of transcripts for the *Atoh1* and *Spdef* genes and low expression of *Tcf7l2* and other members of the Tcf/Lef family (we named the cluster Tcf7l2 “low” cells). Tuft cell precursors expressing doublecortin-like kinase 1 (*Dclk1*) (83), together with tdTomato-positive secretory precursors, formed cluster 12. Immune cells, which were presumably isolated together with epithelial cells (positive for *Lyz2* and various other markers such as *CD14* and immunoglobulin kappa constant, *Igkc*), were clearly delineated as cluster 9. Cell cluster 3, which we labeled "secretory progenitor cells" based on the expression of *Atoh1* (30) was significantly enriched in Defa6-tdTom cells (29% vs. 9% in all epithelial cells isolated from the dissected tumors) (Supplementary Fig. S7B). In addition to *Atoh1*, cells in this cluster also expressed *Lgr5*, *Sox9* and a number of goblet cell markers, such as the aforementioned *Tff3*, *Muc2* and *Agr2*, as well as *Spdef* (84) and *Spink4* (85). The cells in this cluster also expressed genes typical of Paneth cells (e.g., *Lyz1*, *Mmp7*) or colonic enteroendocrine cells (*Reg4*) (86) (Supplementary Table S6). It should be noted that the latter two genes are produced by DCS (see Introduction) and were enriched in Defa6-tdTom tumor cells (Fig. 6D; genes in green).

Next, we investigated the effects of Tcf4 deficiency in tumor cells expressing Defa6-iCre. We crossed *Tcf7l2^flox/flox^* mice with *Apc^+/Min^/ROSA26-tdTomato/Defa6-iCre* mice and analyzed the tumors that developed in the offspring with the *Tcf7l2^flox/flox^/Apc^+/Min^/ROSA26-tdTomato/Defa6-iCre* genotype. Mice with the *Tcf7l2^wt/wt^/Apc^+/Min^/ROSA26-tdTomato/Defa6-iCre* genotype served as controls. In Tcf4-deficient adenomas, we observed a significant decrease in the number of tdTomato-positive glandular structures. The red fluorescent cells were scattered throughout the tumor tissue (Fig. 7A; Supplementary Fig. S8). Subsequent immunohistochemical analysis revealed that the Defa6-tdTom cells in Tcf4 knockout tumors had reduced proliferation, as evidenced by the absence of PCNA protein production. However, Krt20 and Muc2 were detected in these cells, indicating epithelial cell differentiation. It is important to note that strongly Krt20-positive cells were identified within the "red" cells in the glandular structures of wt Tcf4 mice. We also detected co-expression of Muc2 and tdTomato in wt Tcf4 mice, but mainly outside the glandular structures. These observations suggest that the clusters of more differentiated cells, particularly those in clusters 4 and 11 (Fig. 6CD), do not arise from "contaminated" healthy tissues, but are a regular component of tumors.

**Figure 7.**
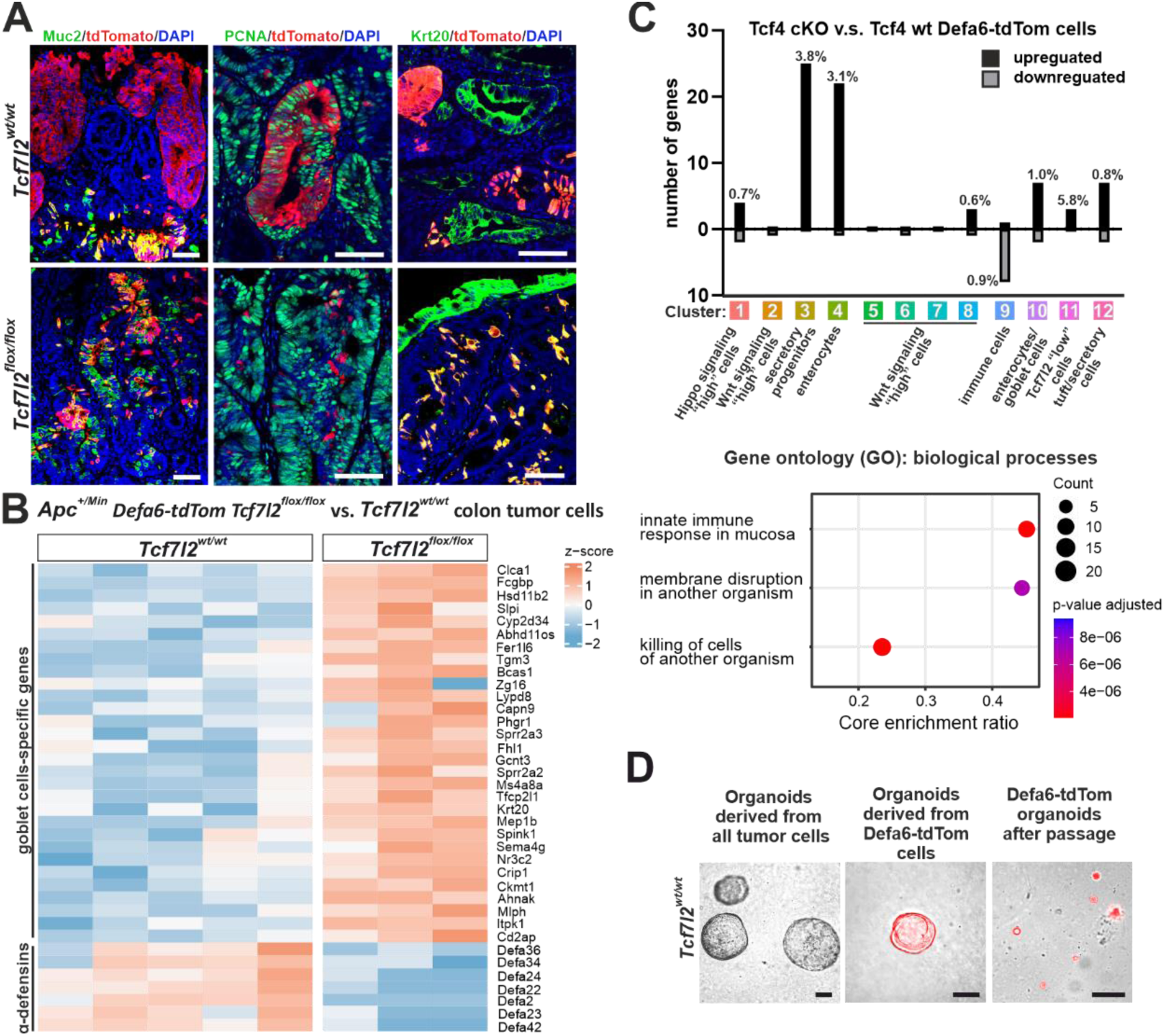
Changes in α-defensin expression and goblet cell phenotype due to disruption of the *Tcf7l2* gene. Mice with a *Tcf7l2^flox/flox^* genotype were crossed with *Apc^+/Min^/ROSA26-tdTomato/Defa6-iCre* mice to generate progeny with Defa6-tdTom cells carrying either wt *Tcf7l2* alleles (*Tcf7l2^wt/wt^*) or a conditional knockout of *Tcf7l2* (*Tcf7l2^cKO/cKO^*). A) Fluorescence microscopy shows co-localization of Defa6-tdTom cells (red signal) with antibodies against Muc2, PCNA and Krt20 (all generating green signal). Scale bar: 0.15 mm. B) Heatmap showing scaled expression obtained by bulk RNA-seq of Defa6-tdTom cells isolated from colon adenomas comparing *Tcf7l2^flox/flox^* to *Tcf7l2^wt/wt^* colon tumor cells. Differential gene expression of genes associated with secretory goblet cells and Reg4^+^ cells is indicated. A complete list of differentially expressed genes can be found in Supplementary Table S7. C) Top, overlaps between differentially expressed genes in Defa6-tdTom tumor cells carrying cKO alleles of the *Tcf7l2* gene (compared to cells with wt *Tcf7l2*) and genes significantly enriched in cellular clusters obtained after scRNA-seq analysis of Defa6-tdTom cells isolated from colorectal tumors. The values next to the selected bars indicate the percentage of genes in each cluster that belong to the differentially expressed genes obtained by bulk RNA-seq. Bottom, GO biological processes enriched in Tcf7l2-deficient Defa6-tdTom cells are depicted, showing the three most affected pathways based on adjusted p-values. D) Representative images of organoid cultures derived from both non-fluorescent total tumor cells and Defa6-tdTom cells isolated from colon tumors of *Tcf7l2^wt/wt^* mice. To promote tumorigenesis, mice were treated with AOM/DSS for 10 weeks prior to cell harvesting. After sorting, cells were cultured in organoid media, encapsulated in Matrigel and cultured in complete organoid culture medium (ENR) containing Wnt surrogate ligand (WntSur). Images were taken after 7 days of initial culture or 4 days after first passage. Scale bar: 0.1 mm.

We next sorted and analyzed Defa6-tdTom tumor cells from the colon tumors of *Apc^+/Min^/ROSA26-tdTomato/Defa6-iCre* and *Tcf7l2^flox/flox^/Apc^+/Min^/ROSA26-tdTomato/Defa6-iCre* mice using bulk RNA-seq. We found downregulation of several α-defensin genes and upregulation of genes predominantly associated with goblet cells (49) (see Fig. 7B for a heatmap representation and Supplementary Table S7 for the full list of differentially expressed genes). When comparing the differentially expressed genes from the bulk RNA-seq with the cellular clusters derived from the scRNA-seq, it was found that the largest overlap among the genes with increased expression after inactivation of the *Tcf7l2* gene was found in clusters 3 and 4. The overlap of genes with decreased expression (after inactivation of *Tcf7l2*) was minimal, unless we used cluster number 9 (immune cells) (Fig. 7C, top). In addition, GSEA revealed that the most altered biological processes were related to "innate immune response in mucosa", "membrane disruption in another organism and antimicrobial response" and "killing of cells of another organism" (Fig. 7C, bottom). It is important to emphasize that these terms were identified after a marked decrease in α-defensin gene expression in Tcf4-deficient tumors. After FACS isolation and Matrigel culture of colon adenoma cells, we confirmed the ability of Defa6-tdTom cells to form organoids. Remarkably, the organoid-forming efficiency of Defa6-tdTom cells was significantly reduced compared to non-tdTomato-labeled tumor cells, with the former forming smaller and slower proliferating organoids after the first passage (Fig. 7D).

## DISCUSSION

### Inactivation of the *Tcf7l2* gene as a model for genetic damage to the intestine

In our study, we used partial (incomplete) inactivation of the *Tcf12* gene to investigate the regeneration process of the intestinal epithelium. In addition to the classical models of epithelial damage by radiation (77, 87, 88), infection (89) or mechanical/chemical damage with DSS (79, 90), we added a model of "genetic damage" and subsequent regeneration. Strikingly, we did not observe activation of the so-called oncofetal (regenerative) program typically associated with activation of genes regulated by the Hippo pathway, such as *Anxa1*, *Ly6a* and *Tacstd2* (77, 79, 90, 91). We considered the possibility that the regeneration response is postponed to an earlier time point after injury, similar to radiation injury (77). However, our analysis within a four-day interval after tamoxifen administration, when only small buds of regenerating crypts are visible (see (18)), showed no specific activation of the regeneration program (Supplementary Fig. S9). In this context, we would also like to mention that staining four days after tamoxifen administration clearly showed the absence of Olfm4 positivity in the crypts of *Tcf7l2^flox/flox^/Villin-CreER^T2^*mice, indicating a loss of stem cells. It was also evident that the apparent uniformity of regenerating crypts seen when proliferating cells were stained with an anti-Ki67 antibody was not as clear with Olfm4 staining, and we observed hyperproliferative crypts/lesions with varying proportions of Olfm4^+^ cells. This proportion was somewhat related to the size (age) of the lesions, with a significant proportion of cells in small hyperproliferative crypts being Olfm4-positive (Supplementary Fig. S9). This then led to a high representation of *Olfm4* transcripts in scRNA-seq. Consequently, the lesion heterogeneity (input sample) influenced the final distribution of cell populations in the regenerating gut.

Treatment with vancomycin restored epithelial integrity and significantly reduced mortality in mice with (partial) Tcf4 inactivation. It is evident that this phenomenon is likely related to the rapid onset of dysbiosis, which we documented by sequential analysis of bacteria colonizing the digestive tract (Fig. 1D). Dysbiosis, defined as an imbalance in the microbial communities colonizing the gut, has profound effects on the intestinal epithelium (reviewed in (92, 93)). However, based on our analyses, we came to the rather surprising conclusion that cells divide in hyperplastic crypts that are phenotypically similar (at the transcriptional level) to their counterparts in healthy, undamaged crypts. A significant difference was the reduced expression of a number of genes encoding defensins detected by bulk RNA-seq. This phenomenon appears to be related to the morphological and cellular organization of the proliferating formations, which lack a clear crypt compartment, and therefore lack Paneth cells and their progenitors.

Another notable observation was the absence (low expression) of *Tcf/Lef* mRNA in some clusters of Mki67-RFP-positive cells. This particularly affected clusters 14 and 9 (Fig. 2B). Within the Defa6-tdTom cells, cluster 11 (Fig. 4B) was also characterized by very low *Tcf7l2* expression; all of these clusters contained early secretory progenitor cells. Interestingly, cluster 11 also showed very low expression of the *Tcf7l2* gene in cells isolated from colorectal tumors. In addition, the expression of other members of the Tcf/Lef family was almost undetectable. It is also interesting that we found the lowest activity of the Wnt signaling pathway in the mentioned cluster (No. 11), measured by the expression of the target genes of this pathway. This finding would imply that in addition to the production of Wnt ligands and other signaling mediators of this pathway (94–96), the level of canonical Wnt signaling in intestinal cells could be modulated by regulating the expression of *Tcf7l2*. This type of regulation, which to our knowledge has not been previously documented, may play a (significant) role, especially in cells at the bottom of the crypt that are likely exposed to Wnt ligands from neighboring cells (Fig. 5AB).

As previously described (10), the loss of Tcf4 leads to the elimination of Paneth cells, accompanied by their (simultaneous) mislocalization outside the crypts on the villi. We also observed this phenomenon when Lyz-positive cells appeared in the small intestinal villi seven days after inactivation of *Tcf7l2* (Fig. 1B). These were probably Paneth cells that had "migrated" from Tcf4-deficient crypts. These cells are no longer dividing, which is why our scRNA-seq analysis of the dividing cells (labeled with Mki67-RFP) could not detect them. The cells labeled with the Defa6-iCre driver (Defa6-tdTom cells) mainly include precursors of Paneth and goblet cells as well as some of the enterocytes. The enteroendocrine cell precursors were not labeled by the Defa6-iCre driver; however, we were able to detect them after labeling with Mki67-RFP, confirming the enteroendocrine precursor cells to be the first to separate during the differentiation of the dividing crypt cells (Fig. 2B) (97).

Our scRNA-seq analysis of Defa6-tdTom cells showed that continuous loss of Tcf4 in the adult intestine leads to an increase in goblet cell progenitors and a decrease in the Paneth cell lineage (analyzed by scRNA-seq). Cells with a so-called intermediate phenotype (production of mucin and markers typical of Paneth cells), which were observed when using the *Villin-CreERT2* driver (10), were possibly not formed (detected). We were only able to detect them in the developing intestine of young mice (P15; Supplementary Fig. S3B). In addition, immunohistochemical staining did not reveal positivity for lysozyme in any cell. Therefore, we conclude that inactivation of Tcf4 leads to cessation of Paneth cell development and one of the ways to lose existing (mature) cells is their migration out of the crypt. However, bulk RNA-seq also revealed that Tcf4-deficient Defa6-tdTom cells produce markers for enterocytes. Does this indicate a different type of cell with a "mixed" phenotype? Probably not. Comparing the expression levels of individual marker genes in the dataset obtained, it becomes clear (Supplementary Table S5) that genes typical of different cell lineages of intestinal epithelial cells are not only typical of a particular cell type, but are also produced by other cell types, albeit at lower levels. Thus, it is not about the *de novo* expression of a specific marker, in this case an enterocyte marker, but about the differences in the expression of a specific gene when comparing Paneth and goblet cell lines. The expression of these genes is hardly detectable in droplet-based scRNA-seq analyses. However, histological staining showed production of a typical enterocyte marker, Alpi, in Tcf4-deficient cells. Recent research has identified a unique subpopulation of goblet cells in the intestinal epithelium, the so-called intercrypt goblet cells (icGCs). These cells are essential for maintaining the mucus barrier in the colon. In contrast to the conventional goblet cells, the icGCs are located on the colon surface between the crypts and secrete a specific type of mucus. A similar (sub)population of goblet cells has been found in the distal part of the small intestine (49). A typical feature of these cells is their common expression of markers for goblet cells and enterocytes. This suggests that some Defa6-tdTom cells may be non-canonical goblet cells. Another intriguing question is why only a small proportion of the goblet cells on the villi are positive for tdTomato. This can be explained by partial recombination of the reporter allele in the pool of progenitor cells that differentiate into goblet cells. Given the constitutive activity of Defa6-iCre and the high efficiency of recombination in Paneth cells, this seems unlikely. Rather, we are inclined to think that the *Defa6-Cre* driver is expressed in a restricted subset of progenitor cells that differentiate into (a subset of) goblet cells; these cells are not dependent on Tcf4 function.

In the small intestine, Wnt ligands play a pivotal role in regulating stem cell maintenance and directing cellular differentiation and proliferation. The production of these ligands primarily occurs in Paneth cells secreting Wnt3, a key ligand that supports the stem cell population and facilitates the stem cell-driven renewal of the intestinal epithelium. Additionally, subepithelial myofibroblasts, located just outside the epithelial cell layer, also contribute to the Wnt ligand pool, particularly Wnt2/2b, thereby supporting the crypt stem cell environment from the underlying mesenchyme (64). Our analysis has shown that Wnt3 is indeed the major ligand produced in epithelial cells. However, the major source of its expression (at the mRNA level) is mainly in secretory progenitor cells, which was confirmed by labeling these cells with the *Defa6-iCre* driver (Fig. 5AB). The fact that no disruption of homeostasis occurs after the loss of Paneth cells in *Tcf7l2^flox/flox^/Defa6-iCre* mice or after blocking the secretion of Wnt ligands from the producing cells by a conditional *Wls* allele did not surprise us in view of the above (extraepithelial sources of Wnt ligands). Similarly, the lack of an observable phenotype when *Defa6-iCre* and *Pdgfra-CreER^T2^* drivers are used simultaneously can be explained by partial recombination of the cKO alleles and thus partial inactivation of the *Wls* gene. Complete recombination does not occur in organoids either, which could explain the survival of epithelial cells in an *in vitro* environment, where there are no extraepithelial sources of Wnt ligands. An alternative explanation could be that there are other, probably progenitor cells that can produce the Wnt ligands. For example, after depletion of Paneth cells with DTA, enteroendocrine cells or tuft cells localized at the crypt base can serve as a source of Wnt ligands (98).

### Expression of Defa6-iCre in colorectal tumors and its use in tumor cell analysis

The *Defa6-iCre* driver was also used for cell labeling and cellular composition analysis of colorectal tumors. We were prompted to perform these experiments because the induced deletion of the floxed allele of the tumor suppressor gene *Apc* leads to an apparent increase in the expression of genes typically found in the epithelium of the small intestine. This signature is particularly striking in the time intervals from 24 to 48 hours after Apc inactivation and appears to be "masked" in later stages by the appearance of tumor-specific markers (19). Subsequent analysis of tumors induced in *Apc^+/Min^* mice by exposure to a mutagen in combination with DSS-induced intestinal damage confirmed the association between the carcinogen (AOM) and the appearance of labeled, i.e., Defa6-tdTom cells. Interestingly, the labeled cells in early tumors did not form contiguous glandular structures but were scattered over different parts of the lesion. At later time intervals, however, we were able to observe fully labeled glandular structures. These structures gradually increased in size with time elapsing between the appearance of the tumor and its analysis (Fig. 6B). A similar staining pattern was observed in tumors that developed spontaneously in *Apc^+/Min^* mice, i.e., mice that were not treated with AOM and DSS. Remarkably, the mere damage to colon tissue never resulted in cell labeling by the *Defa6-iCre* driver (Fig. 6B). This suggests that the appearance of Defa6-tdTom cells is indeed related to cell transformation and not only to tissue damage. How does this pattern of labeled cells arise? Our hypothesis is that during the formation and growth of a tumor triggered by the loss of Apc, a subset of cells activates a secretory program that results in a “scattered cells” pattern. As the tumor grows, some of these cells may adopt a stem-like phenotype that forms the basis for the glandular structures. The observed process of a (stochastic) reversal of the phenotype of Defa6-tdTom cells from a secretory type (see below), or possibly an earlier activation of *Defa6-iCre* expression in a group of so-called tumor stem cells, is consistent with the fact that the number and size of glandular structures in *Apc^+/Min^* mouse tumors are quite variable, but always correlate with the size (age) of the tumor.

The analysis of the tumor composition using scRNA-seq yielded several interesting results. The comparison of all cells isolated from the tumor with Defa6-tdTom cells showed that both samples contained similar cell clusters, which were represented in comparable proportions. An exception was cluster 3 (secretory progenitor cells), which according to immunohistochemical staining for Muc2 (Fig. 7A) consists of Defa6-tdTom cells distributed throughout the tumor. It appears that these cells make up a substantial part of the tumor in the initial growth phase. When these cells are isolated (sorted) from the tumor, their ability to form organoids is limited (Fig. 7D). In the normal digestive tract, Paneth cells are mainly found in the small intestine. However, in various disease states, they can be abnormally present throughout the digestive tract (99). The presence of Paneth cells in colorectal adenomas was first documented over fifty years ago (100). The reported frequencies of the presence of Paneth cells in colorectal adenomas vary considerably, ranging from 0.2% to 39%. Paneth cells are observed more frequently in CRCs than in tubular adenomas (101). A recent study has shown an association between Paneth cell-containing adenomas, male gender and increased adenoma burden (102). In addition, recent research has shown that the accumulation of Paneth cells in early colorectal adenomas is related to β-catenin signaling, suggesting that Paneth cells may form a stem cell niche for adenoma cells, which could have therapeutic implications (103).

### Wnt signaling in tumor cells

Several clusters identified by scRNA-seq analysis showed a high level of Wnt signaling activation. This was expected since the tumor process was triggered by the loss of Apc and subsequent hyperactivation of Wnt signaling. Interestingly, we were able to follow a developmental trajectory. Most clusters with a "high" Wnt signature were directed towards cluster number 5, which consisted of cells with the highest expression of target genes of the Wnt signaling pathway and the highest proliferation. In addition, the expression levels of various Wnt genes within the tumor were remarkable. *Wnt3* was almost exclusively produced in secretory progenitor cells (cluster 3), but it was evident that additional genes encoding Wnt ligands were activated in the terminal cluster 5, particularly the *Wnt6* and *Wnt10b* genes (Fig. 6E). Regarding the signaling pathways involved, Wnt6 has been shown to act via the non-canonical pathway, specifically the planar cell polarity pathway, rather than canonical β-catenin-dependent signaling (104). This shows that different cell types in the tumor influence each other in this way and can also influence cells in the tumor stroma.

Another interesting fact was that the RNA trajectories to cluster 5 contained cells that showed activation of genes regulated by the Hippo signaling pathway. The activation of these genes has been described previously and is particularly associated with intestinal tissue damage in the given model of tumorigenesis (105). Numerous studies have shown that so-called oncofetal genes regulated by the Hippo pathway (or genes indicative of tissue regeneration after injury) are activated to varying degrees in human CRC or in mouse models of tumorigenesis (106). The external signals that trigger such activation and their functional significance are not yet known. Our analysis suggests that the Hippo signature is incorporated into the development of tumor cells, and thus has a dynamic (transient) character.

Finally, inactivation of the *Tcf7l2* gene led to significant suppression of the expression profile of the secretory progenitor cells and to reorientation of these cells towards the goblet cell phenotype (Fig. 7B). Thus, the results obtained with the inactivation of *Tcf7l2* in the small intestine could also be observed to a certain extent in colorectal adenomas. Considering the production of the differentiated epithelial cell marker Krt20 (Fig. 7AB), it could be concluded that a method mimicking the suppression of *Tcf7l2* expression (inhibition of the Wnt pathway) could be a therapeutic approach to the treatment of colorectal tumors. Intriguingly, colorectal cancer characterized by a high number of goblet cells represents a distinct subtype with specific molecular features (107) and different developmental mechanisms compared to non-mucinous carcinomas (108). Mucinous colorectal cancer is often associated with a poorer prognosis than non-mucinous types, particularly in advanced stages (109, 110). A plausible explanation is that the abundant mucin may interfere with the effectiveness of chemotherapy by acting as a barrier to the diffusion of chemotherapeutic drugs (111). However, the origin and role of goblet cells in colorectal cancer remain unclear and warrant further investigation.

## MATERIALS AND METHODS

### Experimental mice

The housing of the mice and the *in vivo* experiments were performed in accordance with the Council Directive of the European Communities of November 24, 1986 (86/609/EEC) and the national and institutional guidelines. The animal care and experimental procedures were approved by the Animal Care Committee of the Institute of Molecular Genetics (approval numbers 63/2019, 08/2020, AVCR 6566/2022 SOV II). *Villin-CreER^T2^*mice (112) were kindly provided by S. Robine (Institut Curie, Centre de Recherche, Paris, France). *Apc^+/Min^* mice (68), *Mki67^RFP^* mice (25), *Pdgfra-CreER^T2^* mice (113), *ROSA26-tdTomato* mice [*B6;129S6-Gt(ROSA)26^Sortm14(CAG-tdTomato)Hze^/J*] (114), *ROSA26-CreER^T^*^2^ mice [*B6.129-Gt(ROSA)26^Sortm1(cre/ERT2)Tyj^/J*] (115) and ROSA-DTA mice [*B6.129P2-Gt(ROSA)26^Sortm1(DTA)Lky^/J*] (63) were purchased from The Jackson Laboratory (Bar Harbor, ME, US). *Tcf7l2f^lox/flox^* mice were obtained from the European Conditional Mouse Mutagenesis Program (EUCOMM; Wellcome Trust Sanger Institute) and have been previously described (18). *Defa6-iCre* (38) and *Wls^flox/flox^* (61) mice have also been described previously. Animals were maintained under specific pathogen-free (SPF) conditions and genotyped according to the provider’s protocols or published protocols.

### Cre-mediated gene recombination, colitis and tumor induction, and antibiotic treatment

Adult mice (8-22 weeks old) producing CreER^T2^ were administered 250 mg/kg tamoxifen (Sigma-Aldrich, St. Louis, MO, USA; 100 mg/mL stock solution in ethanol). The tamoxifen solution was mixed with mineral oil (Sigma-Aldrich) prior to a single administration by gavage. Mice were sacrificed by cervical dislocation at the time points indicated in each experiment after a single administration of the tamoxifen solution. Colon tumors were isolated from 13-19 week old Apc-deficient mice. Inflammatory damage to the colon was induced by 2% (w/v) dextran sulfate sodium (DSS; MP Biomedicals, Irvine, CA, USA; MW36–50 kDa) in drinking water for 5 days. After discontinuation of DSS on day 8 (recovery period after colitis), the colons were removed and analyzed. To ensure inflammation-induced tumorigenesis, mice were injected i.p. with azoxymethane (AOM, 10 mg/kg; Sigma-Aldrich) 7 days prior to DSS administration. The mice were sacrificed 3 weeks after discontinuation of DSS. To ensure sufficient numbers of tdTomato^+^ tumor cells for organoid seeding, we used prolonged AOM/DSS treatment in Apc-deficient mice – 7 days after a single injection of AOM (10 mg/kg), 1% (w /v) DSS was administered for 5 days, which was repeated three times at 14-day intervals. Subsequently, the tumors were processed. To eliminate the gut microbiome in *Tcf7l2^flox/flox^/Villin-CreER^T2^* mice, vancomycin hydrochloride (PHR1732, Sigma-Aldrich; working concentration 500 mg/L) was added to the drinking water 7 days before tamoxifen administration in the indicated experiments; the vancomycin-containing water was changed every 7 days.

### Isolation of intestinal tissues and epithelial cells

For immunohistochemical staining, the intestines were dissected, washed in phosphate-buffered saline (PBS), fixed in 10 % buffered formaldehyde solution (Sigma-Aldrich), embedded in paraffin, sectioned and stained. For organoid cultures and gene expression analysis, the intestinal crypts were isolated from the proximal jejunum of the respective mice. The intestinal tube was cut open lengthwise and the villi were carefully scraped off with a coverslip. The tissue was washed in PBS and incubated in 5 mM EDTA solution in PBS (pH 8; Merck Millipore, Burlington, MA, USA) at 4°C for 30 minutes. The solution was then gently shaken to obtain a suspension of crypts. The crypt suspension was sieved through a 70-μm sieve (Corning, Corning, NY, USA) and centrifuged at 300 × g at 4°C for 5 minutes. For tissue isolation for flow cytometry, the villi were not removed and the whole epithelium obtained after incubation with 5 mM EDTA was used. The epithelium was centrifuged at 300 × g for 5 minutes at 4 °C and resuspended in cleavage medium (serum-free Dulbecco’s Modified Eagle’s Medium; DMEM) with dispase (Thermo Fisher Scientific, Waltham, MA, USA; stock solution 100 mg/ml, diluted 1:300) and DNase I (Thermo Fisher Scientific; working concentration 1 U/ml). The epithelium was incubated 3 x 5 minutes at 37°C on a rotating platform (800 × RPM, 5 minutes, 37°C). Alternatively, colon tumors were harvested directly from the epithelium and cut into small pieces in a cleavage medium with added collagenase type II (C6885, Sigma-Aldrich; working concentration 1 µg/ml). The tumors were incubated 3 × 10 min at 37°C on a rotating platform (800 × RPM, 5 min, 37°C). After each incubation, the tissues were pipetted up and down with a cut tip, and the solution containing the released cells was transferred to DMEM with 10% fetal bovine serum (FCS) to stop cleavage. The collected cells were centrifuged at 300 × g for 5 min at 4 °C and stained.

### Fluorescence-activated cell sorting (FACS)

Epithelial crypt cells from the ileum of *Mki67-RFP Tcf7l2^flox/flox^ Villin-CreER^T2^* mice were stained with Pacific Blue™ (PB) conjugated anti-CD45 antibody (#103126, BioLegend, San Diego, CA, USA; dilution 1:200), PB-conjugated anti-CD31 (#102422, BioLegend; 1:200), fluorescein (FITC)-conjugated anti-EpCAM antibody (#11-5791-82, Thermo Fisher Scientific; 1:400) and allophycocyanin (APC)-conjugated anti CD-24 antibody (#17-0242-82, Thermo Fisher Scientific; 1:400) for 20 min at 4°C; shortly before sorting, Hoechst 33258 (Merck Millipore) was added to the cell suspension. Cells were sorted by forward scatter (FSC), side scatter (SSC), and negative staining for Hoechst and PB. EpCAM^+^ (epithelial) cells were further sorted for RFP and CD24 expression to obtain EpCAM^+^ RFP^+^ CD24^high^ cells, i.e., proliferating epithelial crypt cells. The same staining and sorting strategy was used for *Tcf7l2^flox/flox^/ROSA26-tdTomato/Defa6-iCre* mice; the red fluorescence of tdTomato was used to distinguish recombined cells. Cell sorting was performed using the Influx Cell Sorter (BD Biosciences, San Jose, CA, USA).

### Organoid cultures

Epithelial crypts from resected mouse intestines were embedded in Matrigel (Corning) and cultured as previously described (116). Complete organoid culture medium (ENR): Advanced DMEM/F12 culture medium (Thermo Fisher Scientific) was supplemented with GlutaMax (Thermo Fisher Scientific), 10mM HEPES (1M stock, Thermo Fisher Scientific), penicillin/streptomycin (Thermo Fisher Scientific), B27 Supplement (Thermo Fisher Scientific), N2 Supplement (Thermo Fisher Scientific), 1.25mM N-acetylcysteine (Merck Millipore), 50 μg/ml recombinant mouse epidermal growth factor (EGF; Thermo Fisher Scientific), 2 μl/ml Primocin® (InvivoGen, Toulouse, France) and conditioned culture medium (CM) of mNoggin-Fc (117, 118) and R-Spondin 1 (Rspo1) (117) at a final concentration of 10% CM each. Cells producing the indicated secreted proteins were kindly provided by H. Clevers (Hubrecht Laboratory, Utrecht, Netherlands) and K. Cuo (Stanford University, USA), respectively. If required, an additional 0.5nM Wnt surrogate Fc fusion protein (WntSur; U-Protein Express BV, Utrecht, The Netherlands) was added to the culture medium. Using FACS-sorted tdTomato^+^ cells, 10,000 to 20,000 cells were collected in ENR medium containing the Rho-associated protein kinases (Rock) inhibitor Y-27632 (2.5 mM, Sigma-Aldrich) and 5% Matrigel (Corning). Cells were centrifuged at 300 × g for 5 min at 4 °C, embedded in Matrigel and cultured in ENR medium containing WntSur and Rock inhibitor Y-27632 (2.5 mM) for at least 5 days. After the first passage, WntSur and Y-27632 were removed from the culture medium. Cre-mediated recombination in the organoids was induced by adding 4-hydroxytamoxifen (4-OHT) (Sigma-Aldrich; final concentration 2 µM, 1 mM stock solution was prepared in ethanol) to the culture media. Organoids in culture were imaged using a Leica DMI8 wide-field inverted microscope.

### Immunohistochemical staining and antibodies

A detailed protocol of immunohistochemical staining of paraffin-embedded tissues (19) and organoids (18) has already been described. Primary antibodies: anti-Alpi (rabbit polyclonal, PA5-22210, Thermo Fisher Scientific); anti-ChgA (rabbit polyclonal, ab15160, Abcam, Cambridge, UK); anti-cleaved Casp3 (rabbit monoclonal, #9664, Cell Signaling Technology, Danvers, MA, USA); anti-Krt20 (mouse monoclonal, M7019, Agilent Dako, Santa Clara, CA, USA); anti-Lysozyme (rabbit polyclonal, A0099, Agilent Dako); anti-Muc2 (rabbit polyclonal, sc-15334, Santa Cruz Biotechnology, Dallas, TX, USA); anti-Olfm4 (rabbit monoclonal, #39141, Cell Signaling Technology); anti-PCNA (rabbit polyclonal, ab18197, Abcam); anti-PCNA (mouse monoclonal, ab29, Abcam); anti-Pdgfra (goat polyclonal, AF1062, R&D Systems, Minneapolis, MN, USA); anti-Reg3b (sheep polyclonal, AF5110, R&D Systems); anti-RFP (rabbit polyclonal, 600-401-379, Rockland, Pottstown, PA, USA); anti-RFP (mouse monoclonal, MA5-15257, Thermo Fisher Scientific); anti-Tacstd2 (rabbit monoclonal, ab214488, Abcam); anti-Tcf4 (rabbit monoclonal, #2569, Cell Signaling Technology); anti-Tcf4 (rabbit monoclonal, MA5-35295, Thermo Fisher Scientific). Secondary antibodies (all from Thermo Fisher Scientific): goat anti-rabbit IgG (H+L) Alexa Fluor™ 488 (A11034); goat anti-rabbit IgG (H+L) Alexa Fluor™ Plus 594 (A32740); goat anti-mouse IgG (H+L) Alexa Fluor™ 488 (A11001); goat anti-mouse IgG (H+L) Alexa Fluor™ 594 (A11005); donkey anti-goat IgG (H+L) Alexa Fluor™ 488 (A11055); donkey anti-sheep IgG (H+L) Alexa Fluor™ 488 (A11015). Cells were counterstained with DAPI nuclear stain (Sigma-Aldrich). Microscopic images were acquired with the Leica Stellaris confocal platform or, in the case of organoids, with the Andor Dragonfly 503 Spinning Disk confocal microscope. The images were processed and analyzed with the FiJi package (119).

### DNA and RNA isolation, polymerase chain reaction (PCR), cDNA synthesis and reverse transcription - quantitative PCR (RT-qPCR)

Total RNA from intestinal epithelial cells (freshly isolated crypts or FACS-isolated samples) was isolated using the RNeasy Micro Kit (Qiagen, Germantown, MD, USA) and reverse transcribed using MAXIMA Reverse Transcriptase (Thermo Fisher Scientific) according to the manufacturer’s protocol. RT-qPCR was performed in triplicate using the SYBR Green I Master Mix and the LightCycler 480 instrument (Roche Diagnostics, Indianapolis, IN, USA). For simultaneous isolation of genomic DNA from sorted cells, we used the AllPrep DNA/RNA Micro Kit (Qiagen). The presence of the floxed and recombined allele was analyzed by PCR using the EliZyme™ HS Robust Mix (Elisabeth Pharmacon, Prague, Czech Republic). The primers used for PCR and RT-qPCR are listed in Supplementary Table S1. Isolation and microarray analysis of *Apc^cKO/cKO^/VillinCreER^T2^*colonic epithelium 2 and 4 days after recombination was described in a previous study (19).

### Bulk RNA sequencing (bulk RNA-seq) and computer analysis

The quantity and quality of the isolated RNA was measured with the NanoDrop ND-2000 (NanoDrop Technologies, Wilmington, DE, USA) and analyzed with the Agilent 2100 Bioanalyzer (Agilent Technologies, Santa Clara, CA, USA). The isolated RNA was processed using the Takara Smarter Stranded Total RNA-seq Kit v2 Pico Input Mammalian according to the manufacturer’s instructions. Libraries were sequenced using either the NextSeq 500 or 2000 instrument (both Illumina, CA, USA), with the length set to 75 bases for *Mik67^RFP^/Tcf7l2^flox/flox^/Villin-CreER^T2^*and 122 bases for *Tcf7l2^flox/flox^/ROSA26-tdTomato/Defa6-iCre* libraries. Subsequent processing of the *Mik67^RFP^/Tcf7l2^flox/flox^/Villin-CreER^T2^*data was performed using the nf-core/rnaseq version 1.4.2 bioinformatics pipeline (120). The individual steps included the removal of sequencing adapters and low-quality reads with Trim Galore! (www.bioinformatics.babraham.ac.uk/projects/trim_galore), mapping to the reference genome GRCm38 (Ensembl annotation version 98) (121) with HISAT2 (122) and quantification of gene expression with FeatureCounts (123). The estimated expression per gene served as input for differential expression analysis using the DESeq2 R Bioconductor package (124). Only genes that were expressed in at least two samples were considered for the test. We compared expression between sample groups based on the *Tcf7l2* conditional knock-out (cKO) status of most epithelial cells. Genes that had a minimum absolute log_2_ subject change of 1 (|log_2_ FC| ≥ 1) and statistical significance (adjusted P-value < 0.05) between the compared sample groups were considered differentially expressed.

Data from *Tcf7l2^flox/flox^/ROSA26-tdTomato/Defa6-iCre* libraries were analyzed with the nf-core/rnaseq pipeline (120) version 3.12 using STAR (125) and Salmon (126) to quantify expression per gene using the GRCm39 assembly (Ensembl annotation version 104). Minimal expression (counts) of 10 per gene across samples. Differentially expressed genes were identified using a minimum absolute log_2_ fold change of 1 (|log_2_ FC| ≥ 1) and statistical significance (adjusted P-value < 0.1). In addition, the enrichment of KEGG pathways and Gene Ontologies (GO terms) was analyzed using the GSEA method implemented in the ClusterProfiler R Bioconductor package (127). The initial mapping and analysis of the sequencing data was performed by the Genomics and Bioinformatics Core Facility at the Institute of Molecular Genetics of the Czech Academy of Sciences.

### Single-cell RNA sequencing (scRNA-seq) and computer-assisted analyses

Barcoded single-cell cDNA libraries were prepared with The Chromium Controller (10X Genomics, Pleasanton, CA, USA) using the Chromium Next Gem Single Cell 30 Kit, v3.1, according to the manufacturer’s protocol. The barcoded cDNA was then pooled and sequenced using the NextSeq 500 instrument (Illumina) with an mRNA fragment read length of 130 bases (119 bases for the RFP_WT_GERM sample). We used the 10X Genomics Cell Ranger analysis suite to quantify gene expression per cell based on the GRCm38 Ensembl 98 genome (GRCm39 Ensembl 104 for RFP_WT_GERM). The obtained sequencing data were then analyzed in R Studio with Rx64 3.6.2.Ink (R-tools Technology Inc., Richmond Hill, Canada) using Seurat version 3.1 (128). Only cells with a number of 200 to 7,000 genes and genes detected in more than three cells were included in the quality control. In the experiment analyzing proliferating epithelial cells, cells with more than 10 % mitochondrial genes were excluded from the analysis, resulting in 5,215 cells with 14,308 genes in the *Tcf7l2* wild-type (wt) and 3,859 cells with 14,704 genes in the *Tcf7l2* cKO datasets. In the colon tumor analysis, we excluded cells that had more than 25% mitochondrial genes due to the higher metabolic activity of the tissue (129). This approach resulted in 1,096 cells with 16,715 genes in the whole tumor dataset and 1,406 cells with 14,457 genes in the sorted tumor tdTomato^+^ cell dataset. In both experiments, the datasets were merged, and normalization, scaling and variable gene selection were performed with default settings. Cell clusters were identified using the Louvain approach based on principal component analysis (PCA). Nonlinear dimensionality reduction by Uniform Manifold Approximation and Projection (UMAP) (130) was applied to visualize the low-dimensional embedding of the data and confirm the cluster assignment of the cells.

For the prediction of future gene expression of cells in clusters, we analyzed the scRNA velocity based on the ratio of spliced and unspliced mRNA (31). First, we used the Python package Velocyto 0.17 to calculate separate expression matrices for spliced and nascent mRNA. In the next step, the velocity vectors per cell were calculated using the workflow of the package scVelo 0.2.4 assuming the standard stochastic model. Finally, the velocities were projected onto the UMAP embedding of cell clusters created with the Seurat toolkit.

### Microbiome analysis and data processing

*Mki67^RFP^/Tcf7l2^flox/flox^/Villin-CreER^T2^*mice (7 Cre^+^ and 8 Cre^-^ littermates) were sacrificed 7 days after tamoxifen-induced recombination and the contents were isolated from the ileum of the small intestine. Total DNA was extracted using the ZymoBIOMICS DNA Miniprep Kit (ZYMO Research, Irvine, CA, USA) by repeated bead beating with the FastPrep Homogenizer (MP Biomedicals). The DNA was then quantified using the Qubit dsDNA High Sensitivity Kit (Thermo Fisher Scientific). Samples were processed as technical duplicates to increase the accuracy of the sequencing data. In addition, a ZymoBIOMICS gut microbiome standard (ZYMO Research) was used as a positive control in preparing the sequencing libraries.

The sequencing libraries were prepared using a two-step PCR procedure (131). The first PCR was performed using Kapa HiFi DNA polymerase (Kapa Biosystems, Wilmington, MA, USA) and primers S-D-Bact-0341-b-S-17 (CCTACGGGGGGNGGCWGCAG) and S-D-Bact-0785-a-A-21 (GACTACHVGGGGTATCTAATCC) targeting the V3 and V4 regions of bacterial 16S. The primers contained inline barcodes at the 5’ end and 10-bp tails that were recognized by the second primer pair. Cycling conditions consisted of initial denaturation (95°C, 3 min), followed by 28 cycles of denaturation (98°C, 20 sec), annealing (55°C, 30 sec) and extension (72°C, 30 sec), with final extension (72°C, 5 min). In the second PCR, the unique indices and sequences were added based on the TruSeq adapters. Cycling conditions consisted of initial denaturation (95 °C, 3 min), followed by 12 cycles of denaturation (98 °C, 20 sec), annealing (55 °C, 30 sec) and extension (72 °C, 30 sec) with final extension (72 °C, 5 min). Products were quantified using QIAxcel Advanced Capillary Electrophoresis (Qiagen), and samples within the library were pooled in equal proportions. Libraries were further purified with SPRIselect beads (Beckman Coulter, Brea, CA, USA) and sequenced on the MGI platform at The Genomics Core Facility, CEITEC (Brno, Czech Republic).

Demultiplexing, primer detection and trimming of the sequencing data were performed using Skewer (132). Reads of low quality (expected error rate per paired-end read > 4) were then eliminated. DADA2 (133) was used to denoise the quality-filtered reads and quantify 16S rRNA Amplicon Sequence Variants (ASVs) in each sample. Chimeric ASVs were detected and eliminated using UCHIME (134) and the Silva database (135). Taxonomic assignment of non-chimeric ASVs was performed using the Ribosomal Database Project (RDP) classifier with 80% confidence threshold (136) and the latest version of the Silva database (135). Using Procrustean analysis, we checked the consistency of haplotype composition between identical profiles and retained only the haplotypes that were present in both technical duplicates. We found a high consistency between the technical duplicates. Chloroplasts as well as sequences that could not be assigned to any bacterial strain were considered as food contaminants or sequencing artifacts and excluded from all downstream analyses. The sequences of technical duplicates were pooled for each sample. The ASV abundance matrix (i.e., the number of ASV reads in each sample), the ASV sequences, their taxonomic annotations and phylogeny were merged into a single database together with the metadata of the samples using the phyloseq package (137) in R (R Core Team 2020, Vienna, Austria; http://www.r-project.org/index.html). Taxonomic analysis was performed using the microViz package (138); relative abundances were calculated. Visualization of microbial composition through the iris plot was performed using the centered log ratio (CLR) transformation of taxa at the genus level. PCA was performed to investigate differences in microbiota composition between groups.

### Statistical analysis

Statistical analyses were performed using R, version 3.6.2. Exploratory data analysis was performed for all parameters. Data are presented as mean +-standard deviation (SD; normally distributed data). Survival probabilities for cancer-specific survival were determined using the Kaplan– Meier method and the log-rank test. The results of qRT-PCR analysis and immunofluorescence quantification in FiJi were analyzed using a one-way ANOVA test; a p-value < 0.05 was considered significant. The normality distribution of the data was checked for each set of measurements.

Graphical abstract was created with BioRender.com.

## DATA AVAILIBILITY STATEMENT

The data that support the findings of this study are openly available either in the ArrayExpress or in the Sequence Read Archive of the National Center for Biotechnology Information and in the supplementary material of this article. MIAME-compliant (Minimum Information About a Microarray Experiment) RNA-seq data have been deposited in the ArrayExpress database under the following accession numbers: E-MTAB-6915, E-MTAB-13606, E-MTAB-13730, E-MTAB-13731, E-MTAB-13747, E-MTAB-13752, and E-MTAB-14009. Microbiome analysis sequencing data are available in the European Nucleotide Archive (ENA) (https://www.ebi.ac.uk/ena/browser/home) the under accession number PRJEB74647.

## AUTHOR CONTRIBUTIONS

VKo and LJ contributed to the conception and design of the study and coordinated the experiments. LJ, MS, DH, JO, VKr, SD and TM performed mouse experiments, immunohistochemical studies, organoid experiments and microscopy. JK, KV and MK performed bioinformatic processing and bulk RNA-seq analyses; LJ and LB supported the analysis of scRNA-seq gene expression data. LJ and VKr supervised the breeding of the transgenic mouse strains. MT, SC and KK performed the analysis of the intestinal microbiome. RSB, DF, KB and TV provided mouse strains that were crucial for the manuscript. LJ and MS analyzed the data, completed the results and wrote parts of the manuscript. VKo wrote the final version of the manuscript. All authors read and approved the submitted manuscript.

## ACKNOWLEDGEMENTS

We thank Sarka Takacova for critical reading of the manuscript, Tomas Brabec for discussion on gut immunology, Jakub Kreisinger for assistance with microbiota analysis, Sarka Kocourkova and Zdenek Cimburek for their technical assistance, and Eva Sloncova and Katerina Galuskova for their administrative and technical support. We thank the CF Genomics CEITEC MU, supported by the NCMG research infrastructure, for assistance in obtaining the scientific data presented in this article.

## SUPPLEMENTARY MATERIAL

### Supplementary Figure and Table Legends

**Supplementary Figure S1.**
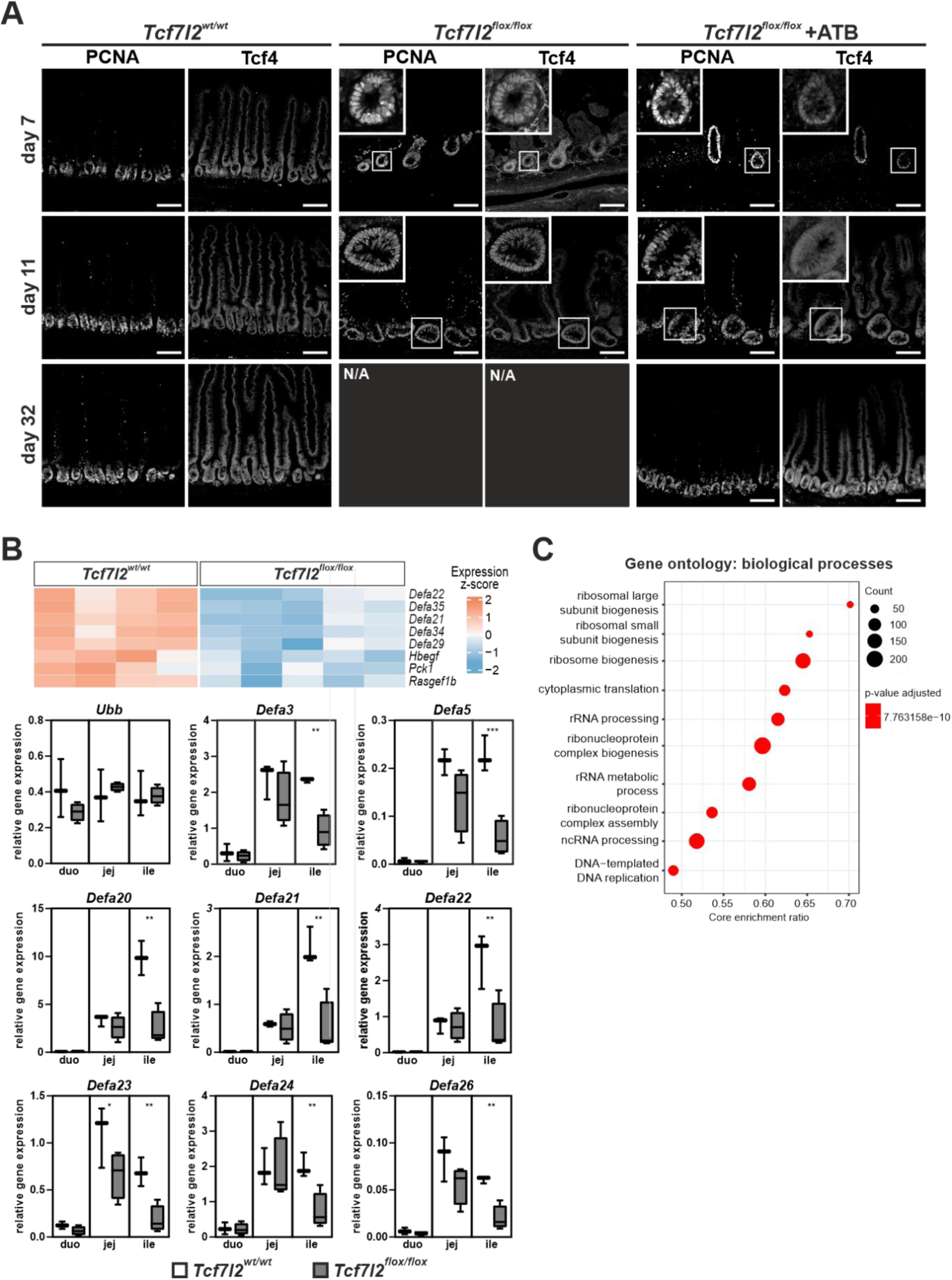
Analysis of the effects of Tcf4 inactivation in the small intestinal epithelium using bulk RNA sequencing (bulk RNA-seq) Mice with the genotype *Tcf7l2^flox/flox^/Mki67^RFP/RFP^/Villin-CreER^T2^*(*Tcf7l2^flox/flox^*) were administered tamoxifen, and proliferating (i.e., Mki67-RFP-positive) epithelial cells were isolated and analyzed after 7 days. Cells from *Tcf7l2^wt/wt^/Mki67^RFP/RFP^/Villin-CreER^T2^*(*Tcf7l2^wt/wt^*) mice served as controls. Hbegf, heparin binding EGF like growth factor; Mki67, proliferation marker Ki-67; Pck1, phosphoenolpyruvate carboxykinase 1; Rasgef1b, RasGEF domain family member 1b; RFP, red fluorescent protein; Tcf7l2, transcription factor 7 like 2; wt, wild-type. A) Grayscale version of the photomicrographs shown in Fig. 1A. B) Top, microarray expression profiling of Mki67-RFP-positive cells. The heatmap shows differentially expressed antimicrobial genes with color scaling indicating the expression level in *Tcf7l2^flox/flox^*mice compared to the *Tcf7l2^wt/wt^* tissue. Significant genes (adjusted p-value < 0.05 and |fold change (FC)| ≥ 2) are shown. Bottom, decreased expression of several α-defensin (*Defa*) genes in *Tcf7l2^flox/flox^*(n = 4) compared to *Tcf7l2^wt/wt^* (n = 3) epithelium was confirmed by reverse transcription and quantitative PCR analysis (RT-qPCR) of isolated crypts from the duodenum (duo), jejunum (jej) and ileum (ile). The phenotype is most pronounced in the distal part of the small intestine. The relative expression of the β-actin gene in the individual intestinal regions was arbitrarily set to 1; the expression of the housekeeping gene ubiquitin B (*Ubb*) is shown in the first graph. The framed areas correspond to the second and third quartiles; the median of the relative expression for each gene is shown as a black line. Statistical significance was determined using the one-way ANOVA test; *p < 0.05; **p < 0.01; ***p < 0.001. C) Gene Set Enrichment Analysis (GSEA) shows the ten most altered biological processes in *Tcf7l2^flox/flox^* cells compared to *Tcf7l2^wt/wt^* cells.

**Supplementary Figure S2.**
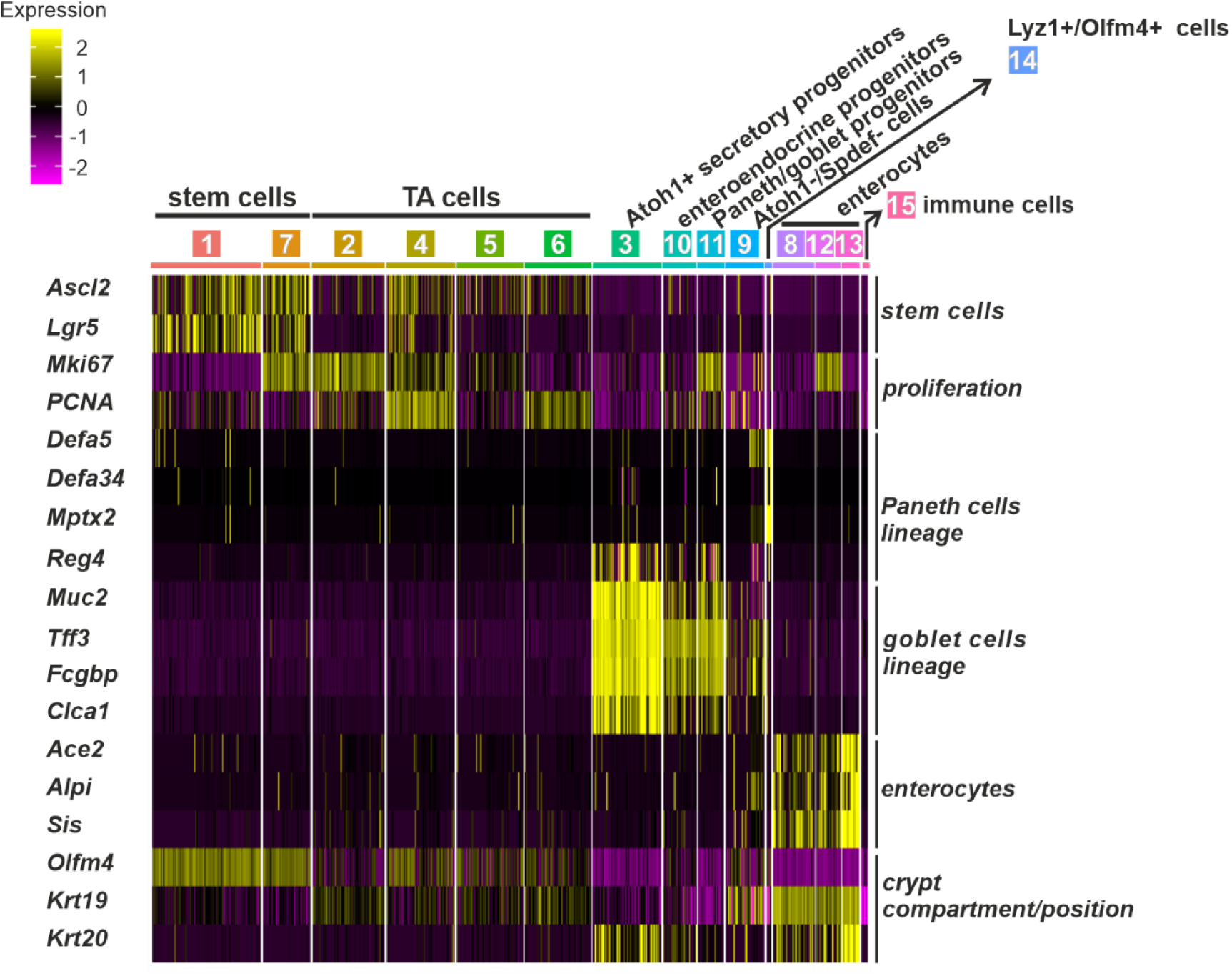
Single cell (sc) RNA-seq analysis of the Mki67-RFP-positive cells isolated from the *Tcf7l2^flox/flox^* and *Tcf7l2^wt/wt^* mice. Heatmap showing scaled expression of indicated marker genes and genes indicating the position of cells within the crypt compartment. Expression levels are color-coded, with yellow indicating high expression and purple indicating low expression.

**Supplementary Figure S3.**
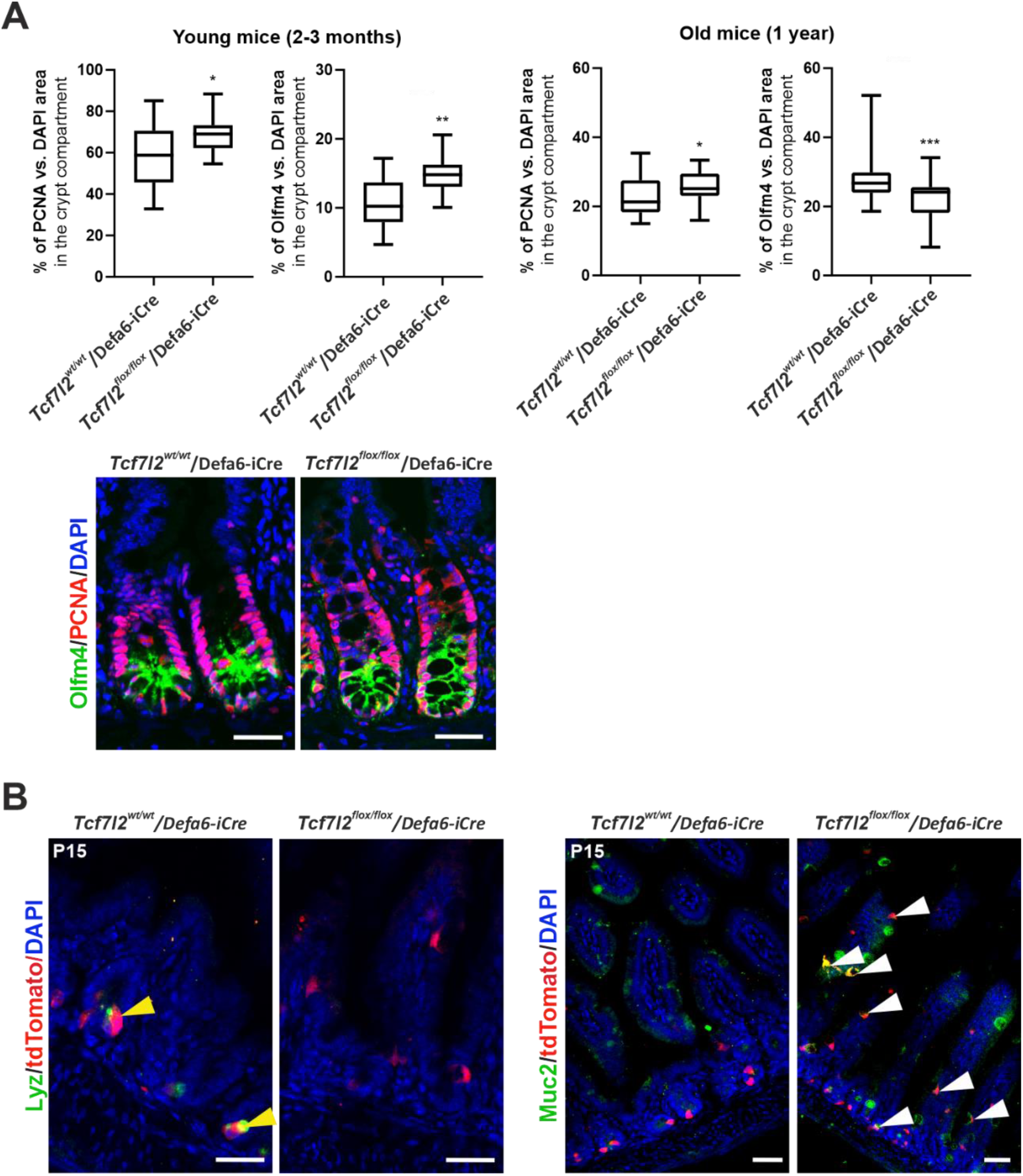
Examination of the morphology of the intestinal epithelium in wt mice (*Tcf7l2^wt/wt^*) and mice carrying the homozygous conditional *Tcf7l2* allele (*Tcf7l2^flox/flox^*) A) Quantification of cells stained positive for anti-proliferating cell nuclear antigen (PCNA) and anti-olfactomedin 4 (Olfm4) in young (2-3 months) and older (1 year) mice of the indicated genotype. For each group, at least ten microscopic fields from three biological replicates were analyzed. The signal of the antibody was normalized to the DAPI counterstain. Statistical significance was determined using the one-way ANOVA test; *p < 0.05; **p < 0.01; ***p < 0.001. Below is a representative image of immunohistochemical staining of PCNA (red nuclear signal) and Olfm4 (green signal) in a young adult mouse intestine. B) Histological slides showing the localization of Defa-tdTom cells (tdTomato) and their colocalization with lysozyme (Lyz; yellow arrowheads) or mucin 2 (Muc2; white arrowheads) in the indicated mouse strains 15 days after birth (P15). Animals were crossed with *Rosa26-tdTomato* reporter mice. Scale bar: 50 μm.

**Supplementary Figure S4.**
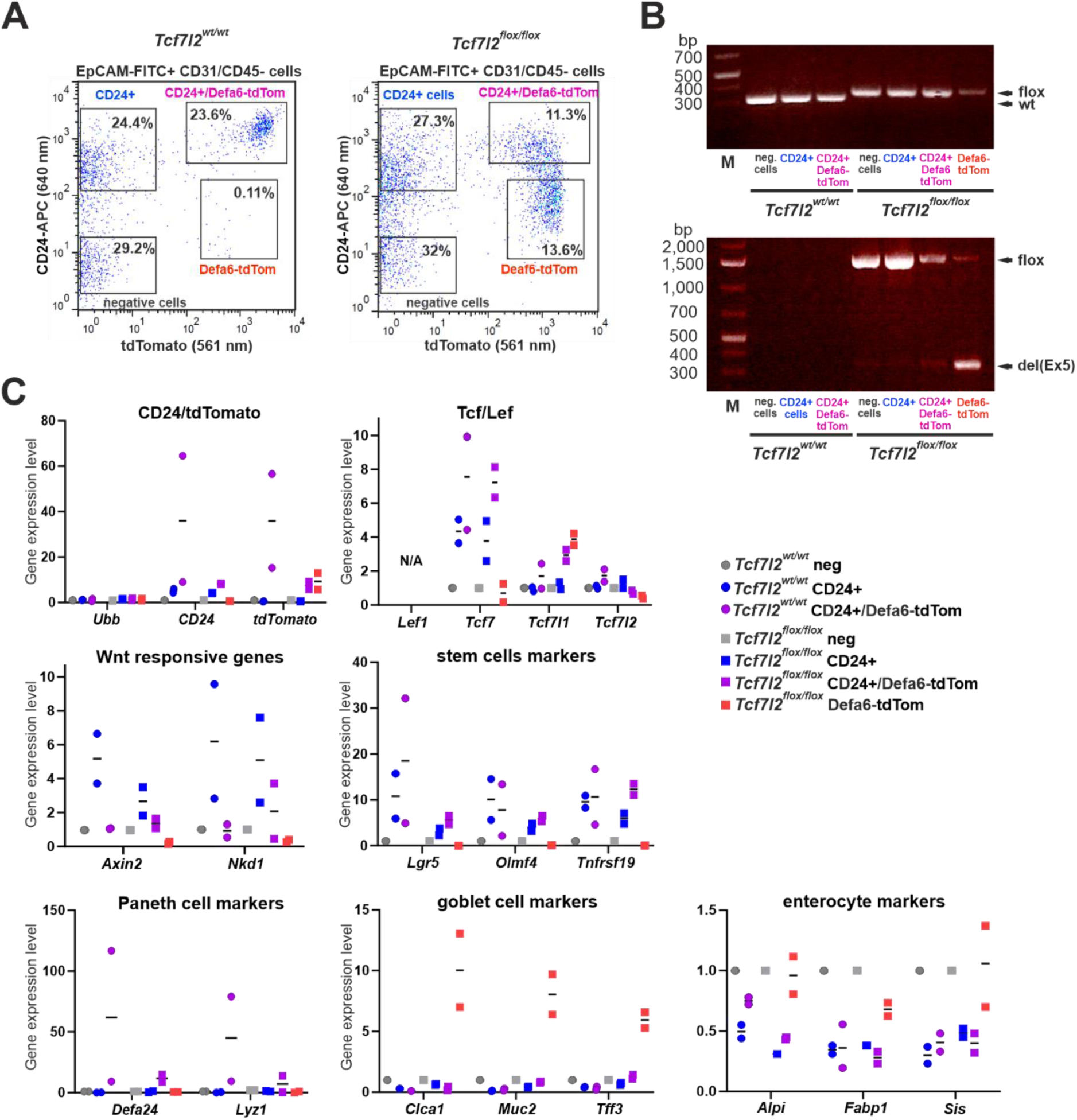
Properties Tcf7l2 wt (*Tcf7l2^wt/wt^*) and Tcf7l2-deficient (*Tcf7l2^flox/flox^*) cells isolated from the intestines of mice of the indicated genotype. The mice were crossed with *Defa6-iCre/Rosa26-tdTomato* mice. A) Representative flow cytometry plots show the gating strategy used to isolate CD24-positive and CD24-negative tdTomato^+^ (Defa6-tdTom) cells. Single cells were harvested from the middle part of the small intestine; viable (Hoechst 33258-negative), epithelial (EpCAM^+^), CD31/CD45-negative cells were sorted based on the surface marker CD24 and endogenous tdTomato fluorescence. EpCAM, epithelial cell adhesion molecule. Percentage of cells in the particular sorting gate is indicated. B) Genotyping of the *Tcf7l2* wt and floxed (flox) alleles (top) and the recombinant Tcf7l2*^del(Ex5)^* allele [del(Ex5); bottom] was performed using DNA from all FACS-isolated cell populations. FACS, fluorescent activated cell sorting. C) Analysis of indicated cell type-specific genes was performed using RNA from the FACS-isolated cell populations by RT-qPCR. The relative gene expression in CD24/tdTomato-negative cells was arbitrarily set to 1. The expression of the housekeeping gene *Ubb* is shown in the first graph. The black lines indicate the mean value for two biological replicates; each reaction was performed in technical triplicates. Alpi, alkaline phosphatase; Clca1, chloride channel accessory 1; Defa24, defensin alpha 24; Fabp1, fatty acid-binding protein 1; Lef1, lymphoid enhancer-binding factor 1; Lgr5, leucine-rich repeat-containing G protein-coupled receptor 5; Nkd1, naked cuticle 1; Sis, sucrase-isomaltase; Tcf7, transcription factor 7; Tcf7l1, transcription factor 7-like 1; Tff3, trefoil factor 3; Tnfrsf19, tumor necrosis factor receptor superfamily member 19.

**Supplementary Figure S5.**
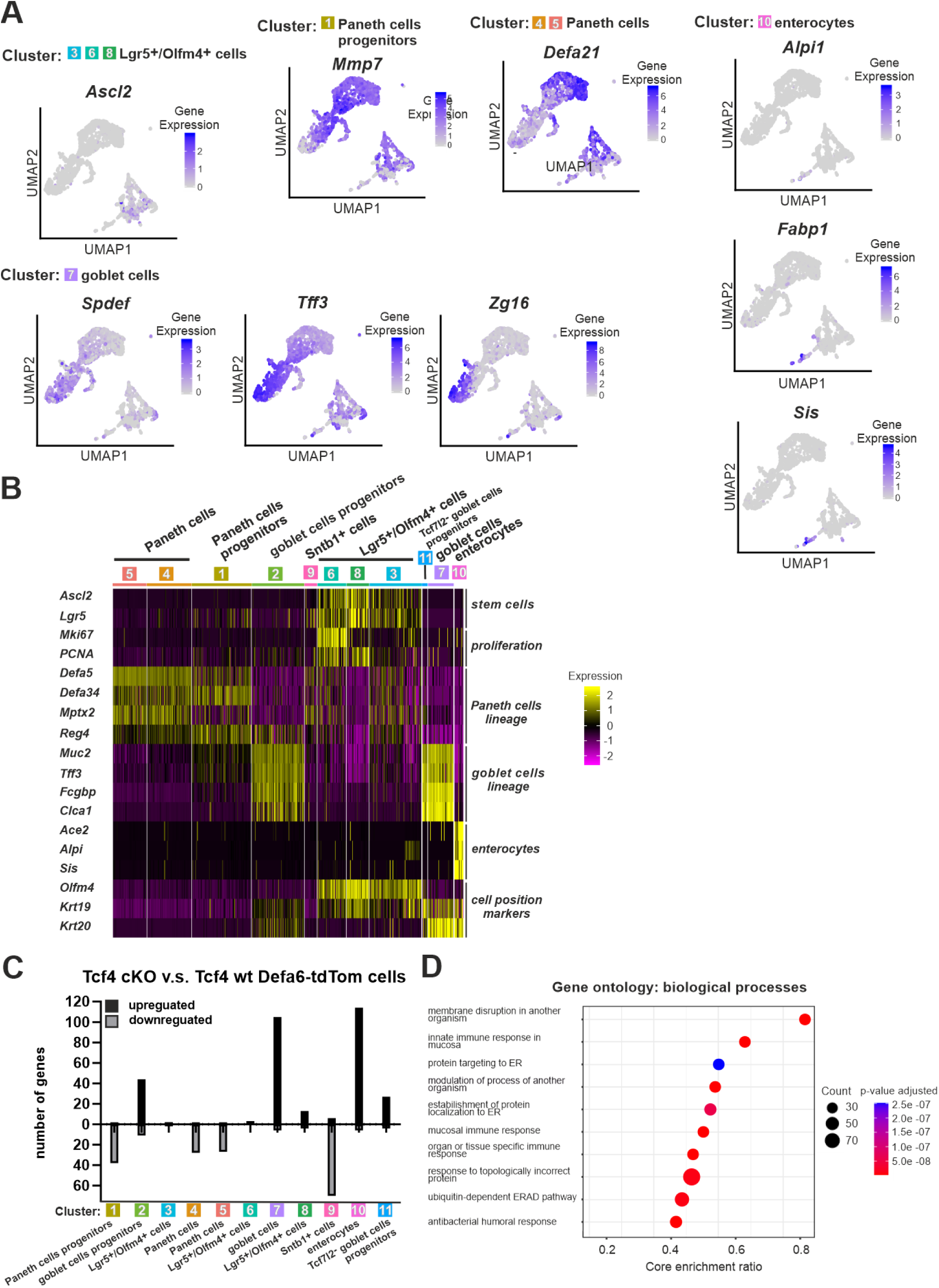
Analysis of the effects of Tcf4 inactivation in Defa6-tdTom cells using scRNA (A) and bulk RNA sequencing (B). The results complement the data shown in Fig. 4. A) UMAP plots showing the expression patterns of the different marker genes. The plots were derived from the combined sample. B) Heatmap showing scaled expression of indicated marker genes and genes indicating the position of cells within the crypt compartment. Expression levels are color-coded, with yellow indicating high expression and purple indicating low expression. Ascl2, achaete-scute family bHLH transcription factor 2; Mmp7, matrix metallopeptidase 7; Spdef, SAM pointed domain containing ETS transcription factor; Zg16, zymogen granule protein 16. C) Overlaps between differentially expressed genes in Defa6-tdTom cells carrying cKO alleles of *Tcf7l2* (compared to cells with wt Tcf7l2) and genes significantly enriched in cellular clusters obtained after scRNA-seq analysis of Defa6-tdTom cells isolated from the small intestinal epithelium. The values next to the selected bars indicate the percentage of genes in each cluster that belong to the differentially expressed genes obtained by bulk RNA-seq. D) Gene Ontology (GO) biological processes enriched in *Tcf7l2*-deficient Defa6-tdTom cells when compared to *Tcf7l2* wt cells. The 10 most affected pathways based on adjusted p-values are depicted. ERAD, endoplasmic reticulum-associated degradation.

**Supplementary Figure S6.**
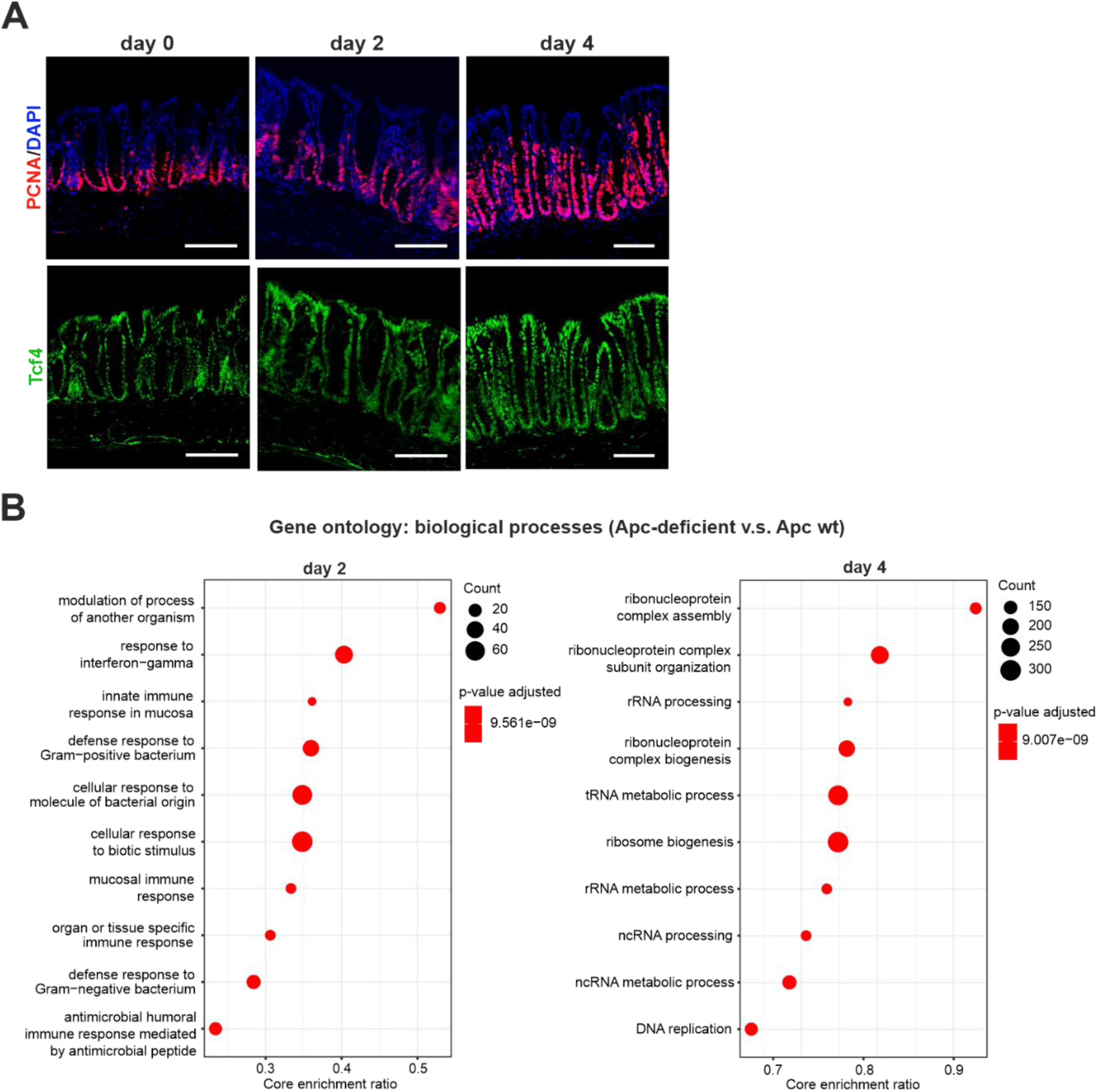
Examination of colon epithelium 2 and 4 days upon tamoxifen-mediated conditional knockout of adenomatous polyposis coli (*Apc^cKO/cKO^*) gene using V*illin-CreER^T2^* recombination driver. A) Representative specimens showing cells stained with the antibody against PCNA (red; upper panel) and Tcf4 (green; lower panel) in the control (day 0) and Apc-deficient (day 2, 4) colon crypts. Note increased proliferation and extension of the crypts 4 days upon Apc truncation. Scale bar: 0.1 mm. B) GSEA showing the 10 most significantly altered biological processes in Apc-deficient colonic crypts compared to control colonic epithelium.

**Supplementary Figure S7.**
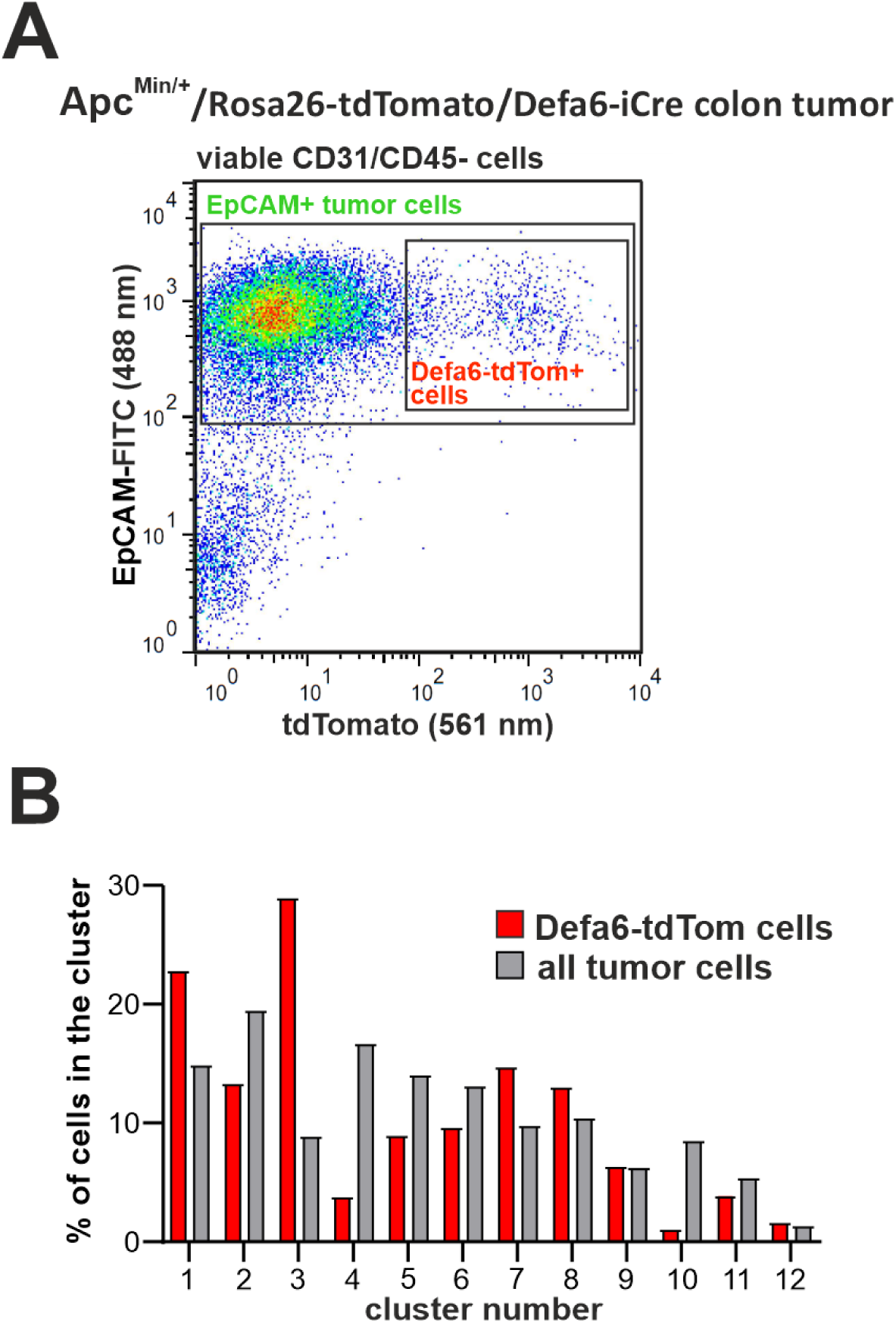
Sorting strategy and abundance of cell clusters identified in the scRNA-seq analysis in the colon adenoma cells. A) Representative diagram showing the fluorescence-activated cell sorting (FACS) strategy for obtaining colon tumor cells from mice with multiple intestinal neoplasia (Min) in the intestinal epithelium. Cells from dissected colonic adenomas from *Apc^Min/+^/Rosa26-tdTomato/Defa6-iCre* mice were sorted for viability (i.e., Hoechst 33258-negative cells) and EpCAM positivity; CD31-positive endothelial cells and CD45-positive leukocytes were excluded from the sorted populations. The Defa6-tdTom cells were distinguished by the red fluorescence of the tdTomato protein (Defa6-tdTom cells). B) Bar graph showing the percentage of each cluster of Defa6-tdTom tumor cells (red) and all cells isolated from the colon tumor (gray).

**Supplementary Figure S8.**
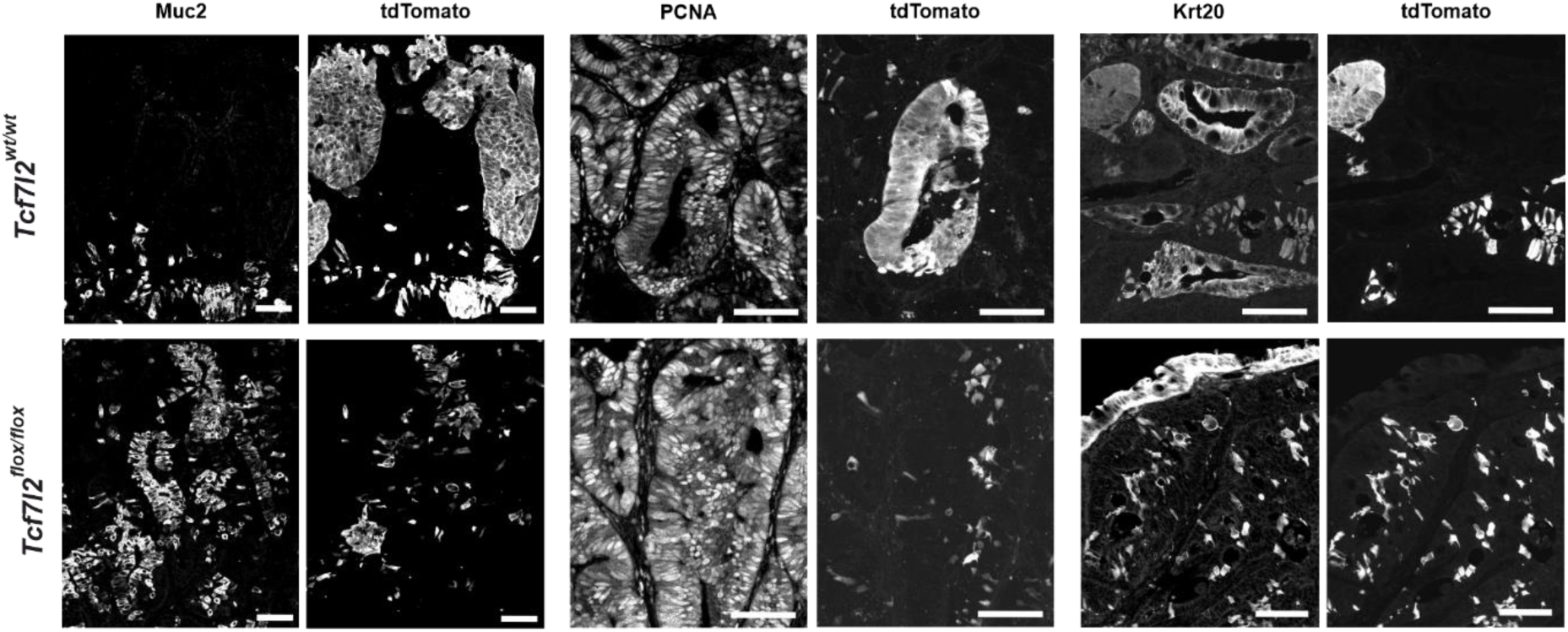
Grayscale version of the photomicrographs shown in Fig. 8A.

**Supplementary Figure S9.**
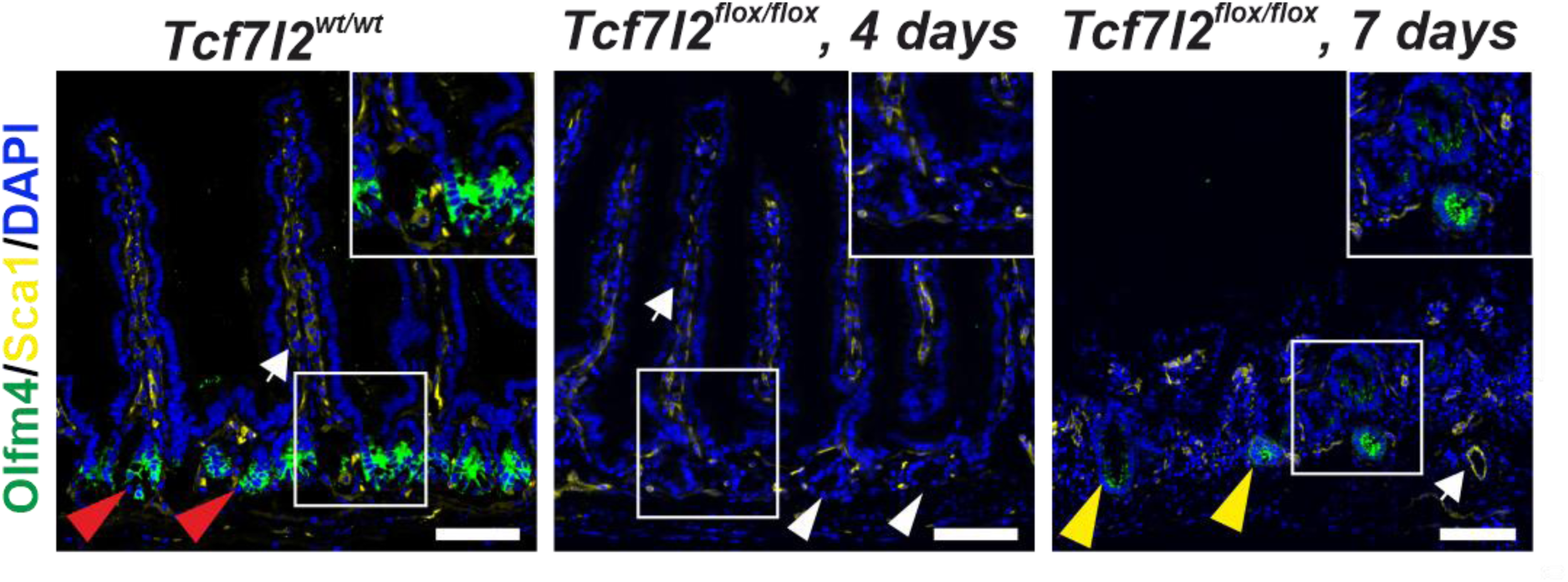
Tcf7l2 knockout effects on small intestinal crypts and Olfm4 dynamics. Histologic analysis shows dynamic changes in the expression of Olfm4 (green fluorescence) and stem cell antigen 1 (Sca1; yellow). White arrows indicate that Sca1 expression is localized at the endothelial surface in mesenchymal tissue. Red arrowheads indicate Olfm4-positive cells in the crypts of *Tcf7l2^wt/wt^* mice; white arrowheads indicate the absence of Olfm4-positive cells and marked deformation of the crypt compartment 4 days after *Tcf7l2* knockout. Remarkably, Olfm4-positive cells appear in the regenerating crypt compartments 7 days after *Tcf7l2* inactivation. Scale bar: 100 μm.

**Supplementary Table S1**

Sequences of primers used in the genotyping PCR and RT-qPCR experiments.

**Supplementary Table S2**

DEGs identified between Mki67-RFP-positive cells obtained from *Tcf7l2* cKO and wt small intestinal crypts. Genes with an adjusted p-value < 0.05 and |log_2_ FC | ≥ 1 were considered significant. False discovery rate (FDR) correction based on the total number of genes in the dataset was used to adjust the p-value. FC, fold change.

**Supplementary Table S3**

The genes specifically expressed in each cluster identified in Mki67-RFP-positive cells in the small intestinal crypts are listed in the corresponding sheet. Genes with the average expression |log_2_ FC| > 0.25 and p-value < 0.05 were considered as significant. FDR correction based on the total number of genes in the dataset was used to adjust the p-value. The percentage of cells with the gene expression in the first and second most significant groups is indicated.

**Supplementary Table S4**

DEGs identified between Defa6-tdTom-positive *Tcf7l2* cKO and wt cells obtained from the ileum of the small intestine. Genes with an adjusted p-value < 0.05 and |log_2_ FC | ≥ 1 were considered significant. FDR correction based on the total number of genes in the dataset was used to adjust the p-value.

**Supplementary Table S5**

The genes specifically expressed in each cluster of Defa6-tdTom^+^ cells in the middle small intestinal crypts are listed in the corresponding sheet. Genes with the average expression |log_2_ FC| > 0.25 and p-value < 0.05 were considered as significant. FDR correction based on the total number of genes in the dataset was used to adjust the p-value. The percentage of cells with the gene expression in the first and second most significant groups is indicated.

**Supplementary Table S6**

The genes specifically expressed in each cluster identified in cells isolated from the colon tumor of *Apc^+/Min^* mice are listed in the corresponding sheet. Genes with the average expression |log_2_ FC | > 0.25 and p-value < 0.05 were considered as significant. Bonferroni correction based on the total number of genes in the dataset was used to calculate the p-value. The percentage of cells with the gene expression in the first and second most significant group is indicated.

**Supplementary Table S7**

Differentially expressed genes (DEGs) between tdTomato-positive *Tcf7l2* cKO and wt cells obtained from the colon tumor of *Apc^+/Min^* mice. Genes with an adjusted p-value < 0.05 and |log_2_ FC | ≥ 0.5 were considered significant. FDR correction based on the total number of genes in the dataset was used to adjust the p-value. Genes characteristic of secretory precursors (pink color) and (blue color) are indicated.

